# Reversal of the renal hyperglycemic memory by targeting sustained tubular p21 expression

**DOI:** 10.1101/2021.07.05.450846

**Authors:** Moh’d Mohanad Al-Dabet, Khurrum Shahzad, Ahmed Elwakiel, Alba Sulaj, Stefan Kopf, Fabian Bock, Ihsan Gadi, Silke Zimmermann, Rajiv Rana, Shruthi Krishnan, Dheerendra Gupta, Sumra Nazir, Ronny Baber, Markus Scholz, Robert Geffers, Peter Rene Mertens, Peter P. Nawroth, John Griffin, Chris Dockendorff, Shrey Kohli, Berend Isermann

## Abstract

A major therapeutic obstacle in diabetes mellitus is the metabolic or hyperglycemic memory: the persistence of impaired organ function despite improvement of blood glucose. Therapies reversing the hyperglycemic memory and thus improving already established organ-dysfunction are lacking, but urgently needed considering the increasing prevalence of diabetes mellitus worldwide. Here we show that glucose-mediated changes in gene expression largely persist in diabetic kidney disease (DKD) despite reversing hyperglycemia. The senescence-associated cyclin-dependent kinase inhibitor p21 (*Cdkn1a*) was the top hit among genes persistently induced by hyperglycemia and was associated with sustained induction of the p53-p21 pathway. Persistent p21 induction was confirmed in various animal models, in several independent human samples and in *in vitro* models. Tubular p21 expression and urinary p21-levels were associated with DKD severity and remained elevated despite improved blood glucose levels in humans, suggesting that p21 may be a biomarker indicating persistent (“memorized”) kidney damage. Glucose-mediated p21 induction and tubular senescence were enhanced in mice with reduced levels of the disease resolving protease activated protein C (aPC). Mechanistically, glucose-induced and sustained tubular p21 expression is linked with demethylation of its promoter and reduced DNMT1 expression. aPC reverses already established p21 expression independent of its anticoagulant function through receptor signaling. Accordingly, new pharmacological approaches specifically mimicking aPC signaling (3K3A-aPC, parmodulin-2) enabled the reversal of glucose-mediated sustained tubular p21 expression, tubular senescence, and DKD. Thus, p21-dependent tubular senescence contributes to the hyperglycemic memory but can be therapeutically targeted.

**Single sentence summary:** aPC signaling targets persistent p21 expression and tubular senescence and reverses the hyperglycemic memory in diabetic kidney disease.

## Introduction

Diabetic kidney disease (DKD) is now the leading cause of end-stage renal disease in industrialized countries, affecting approximately 40% of diabetic patients (*1*). Current therapies, particularly blood sugar control and renin angiotensin aldosterone system (RAAS) inhibition, at best delay disease progression (*2*). Sodium-glucose co-transporter-2 inhibitors (SGLT2i) are promising therapeutic adjuvants, but the long-term consequences, the associated risk profile and their ability to halt or even reverse disease progression remain largely unknown (*3*). The persistence of diabetes-associated complications despite improved blood glucose control is known as metabolic or hyperglycemic memory (*4, 5*). Disease reversal, e.g., the improvement of already established DKD, remains an unmet medical need(*2*). Accumulating evidence suggests that epigenetic mechanisms such as DNA methylation, posttranslational histone modification, and noncoding RNAs are mechanistically linked to hyperglycemic memory (*5, 6*), but these insights have not yet led to new diagnostic or therapeutic approaches.

Several pathomechanisms have been linked with DKD (*7*), yet their potential role in hyperglycemic memory is unknown. The redox regulator p66^Shc^ is epigenetically induced in diabetic patients and mice, and p66^Shc^ deficiency protects against diabetic glomerulopathy in mice (*8-10*). The cytoprotective and signaling-competent coagulation protease-activated protein C (aPC) reverses epigenetically induced p66^Shc^ expression in experimental diabetes (*9, 11*). These data illustrate that (i) epigenetic changes in diabetic vascular disease are in principle reversible and (ii) that the hyperglycemic memory is linked to endothelial dysfunction, as aPC’s plasma levels reflect endothelial function (*12, 13*).

Both hypo- and hypermethylation have been reported in association with diabetes and glomerular and tubulointerstitial damage in the context of DKD (*5, 14, 15*). Tubulointerstitial damage in DKD is characterized by the accumulation of extracellular matrix (fibrosis), inflammation, and senescence of tubular cells (*16-20*). Senescence, a failure of cells to re-enter the cell cycle, may impede tissue repair (*21-23*), has been linked with chronic kidney disease, but its contribution to hyperglycemic memory remains unknown.

Here, we combined hypothesis-free approaches with analyses of human samples, genetic and interventional mouse models and *in vitro* studies to gain insights into mechanisms of hyperglycemic memory. We identified sustained p21 expression as a regulator of hyperglycemic memory. Inhibition of sustained p21 expression erases hyperglycemic memory and improves already established experimental DKD.

## Results

### Identification of genes and pathways associated with hyperglycemic memory

To identify regulators of renal hyperglycemic memory, we used a mouse model of hyperglycemia reversal and conducted hypothesis-free expression analyses (RNAseq). Persistent hyperglycemia was induced in mice for 16 weeks, at which stage mice had developed albuminuria (**Fig. 1a-c**). Subsequently, blood glucose levels were markedly reduced in a subgroup of mice for additional 6 weeks using an SGLT2 inhibitor (SGLT2i, Dapagliflozin®, DM+SGLT2i), while in another subgroup blood glucose levels remained elevated for the entire 22 weeks period (DM-22, **Fig. 1a**,**b**). Reduction of blood glucose levels halted kidney function decline (DM-16 *versus* DM+SGLT2i), resulting in lower albuminuria compared to untreated diabetic mice (DM-22), but albuminuria remained increased compared to non-diabetic control mice (C, **Fig. 1c**). Thus, lowering blood glucose levels failed to reverse already established albuminuria in mice, mimicking the hyperglycemic memory.

**Figure 1:**
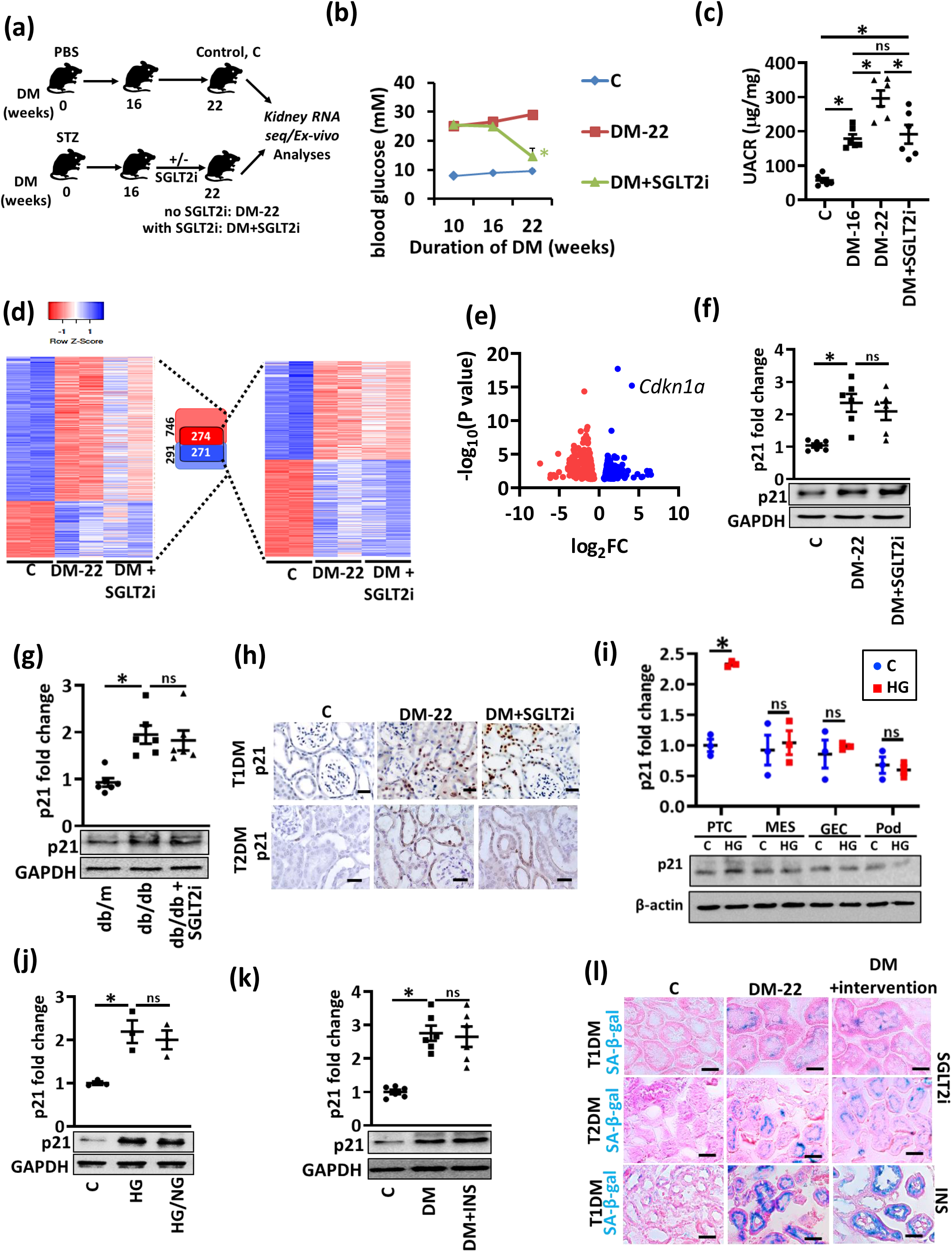
Identification of genes and pathways associated with hyperglycemic memory. **a)** Experimental scheme showing non-diabetic control (C) or diabetic mice without (DM-22) or with intervention to reduce blood glucose levels by sodium/glucose cotransporter-2 inhibitor (Dapagliflozin®, DM+SGLT2i). Mice were age matched and SGLT2i treatment was started after 16 weeks of persistent hyperglycemia (streptozotocin; STZ-induced hyperglycemia) **b)** Average blood glucose levels in the experimental groups (as described in a) after 10 or 16 weeks of persistent hyperglycemia and at 22 weeks. Line graphs reflecting mean±SEM of at least 6 mice per group; ANOVA, \**P*<0.05 **c)** Dot plot summarizing albuminuria (urinary albumin-creatinine ratio, µg albumin/mg creatinine; UACR) in non-diabetic (C) or diabetic mice without (DM-16 and DM-22) or with intervention to reduce blood glucose levels (DM+SGLT2i). Dot plot reflecting mean±SEM of at least 6 mice per group; ANOVA, \**P*<0.05; ns: non-significant **d)** Heat map summarizing gene-expression in non-diabetic control (C) or diabetic mice without (DM-22) or with intervention to reduce blood glucose levels (DM+SGLT2i). All genes with significantly changed expression between control and diabetic mice are shown on the left side of the panel and are illustrated by lightly colored boxes in the middle (number of genes shown in black). The subset of genes with persistently changed expression despite reduced blood glucose levels are shown on the right side of the panel and illustrated by the darker colored smaller boxes in the middle (number of genes shown in white) **e)** Volcano plot summarizing the persistently induced and repressed genes in diabetic mice after blood glucose reduction based on Log FC values and FDR. *Ckdn1a* (p21) belongs to the top persistently induced genes **f**,**g)** Sustained p21 expression *in vivo* (STZ-model, f; db/db model, g) despite reducing glucose levels (DM+SGLT2i, f; db/db+SGLT2i, g) as compared to mice with persistently elevated glucose levels (DM-22, f; db/db, g) or normoglycemic controls (C, f; db/m, g). Exemplary immunoblots, GAPDH: loading control (bottom) and dot plots summarizing results (top). Dot plots reflecting mean±SEM of at least 6 mice per group; ANOVA, \**P*<0.05; ns: non-significant **h)** Hyperglycemia-induced (DM-22; T1DM, STZ-model, top; T2DM, db/db-model, bottom) p21 expression is predominately seen in tubular cells and remains high despite lowering blood glucose levels (DM+SGLT2i). In normoglycemic controls (C) p21 is barely detectable. Exemplary immunohistochemical images, p21 is detected by HRP-DAB reaction, brown; hematoxylin nuclear counter stain, blue. Scale bars represent 20 µm **i)** Exemplary immunoblot loading control: β-actin, bottom) and dot plot summarizing results (top) of p21 () in mouse primary tubular cells (PTC), mouse mesangial cells (MES), mouse glomerular endothelial cells (GEC), or mouse podocytes (Pod) maintained under normo-(5 mM; C) or hyerglycemic (25 mM; HG) conditions. Dot plot reflecting mean±SEM of at least 3 independent experiments; t-test comparing C versus HG for each cell line, \**P*<0.05; ns: non-significant **j)** Sustained p21 expression *in vitro* (Human kidney cells, HEK293) despite reducing glucose levels (HG/NG) as compared to cells with persistently elevated glucose levels (HG) or normoglycemic controls (C). HEK-293 cells were maintained under normal glucose (5 mM, C), high glucose (48 h of 25 mM, HG), or high glucose followed by normal glucose (25 and 5 mM, each for 24 h, HG/NG). Exemplary immunoblots (bottom, GAPDH: loading control) and dot plots summarizing results (top). Dot plots reflecting mean±SEM of 3 independent experiments; ANOVA, \**P*<0.05; ns: non-significant **k)** Sustained p21 expression *in vivo* (STZ-model) despite reducing glucose levels using insulin (DM+INS, j) as compared to mice with persistently elevated glucose levels (DM-22) and normoglycemic controls (C). Exemplary immunoblots (bottom, GAPDH: loading control) and dot plots summarizing results (top). Dot plots reflecting mean±SEM of at least 6 mice per groups; ANOVA, \**P*<0.05; ns: non-significant.**l)** Exemplary histological images of SA-β-gal stain (senescence associated β-galactosidase, blue; eosin counterstain) in T1DM (STZ-model, top and bottom) and T2DM (db/db model, middle) despite reducing glucose levels (DM+intervention; SGLT2i, top and middle or insulin, INS, bottom) as compared to mice with persistently elevated glucose levels (DM-22) or normoglycemic controls (C). Scale bars represent 20 µm.

To identify persistently altered gene expression in association with sustained albuminuria, RNAseq data were compared among nondiabetic control (C), untreated (DM-22) and SGLT2i-treated (DM+SGLT2i) diabetic mice. Compared to controls (C), 291 genes were induced, and 746 genes were suppressed in diabetic mice (DM-22, **Fig. 1d**). Despite reducing blood glucose levels in the SGLT2i-treated mice (DM+SGLT2i), the expression of 271 (out of 291) induced and 274 (out of 746) suppressed genes remained increased or suppressed, respectively, reflecting persistent gene expression possibly contributing to hyperglycemic memory (**Fig. 1d**). The cell-cycle and senescence associated gene *Cdkn1a* (p21) was among the top persistently induced genes (**Fig. 1 e**). Functional annotation of these memorized genes revealed pathways related to metabolism (metabolic and endocrine pathways), growth factor signaling, chemokine and cytokine-mediated inflammation, and p53-p21 pathway and feedback loops as the most prevalent changes (**Supplementary Fig. 1a**). Persistent p21 induction was confirmed at mRNA (**Supplementary Fig. 1b**) and protein level (**Fig. 1f**).

### Persistent induction of tubular p21 expression and senescence in experimental DKD

Induction of p21 in DKD has been reported, but its contribution to hyperglycemic memory has not been shown hitherto (*24*). Using a model of type 2 diabetes mellitus (T2DM; db/db mice) and a similar experimental set up (persistent hyperglycemia until age 16 weeks followed by reduction of blood glucose levels using SGLT2i; **Supplementary Fig. 1c**,**d)**, we likewise observed sustained albuminuria (**Supplementary Fig. 1e**) in association with persistent p21 induction in renal cortex extracts (**Fig. 1g, Supplementary Fig. 1f**).

Immunohistological analyses revealed persistently increased p21 expression despite lowering blood glucose levels in tubular cells in the T1DM and the T2DM models (**Fig. 1h, Supplementary Fig. 2a**,**b**). Predominant p21 induction in tubular cells by high glucose (HG) was confirmed *in vitro* comparing mouse primary tubular cells (PTCs), mesangial cells (MESs), glomerular endothelial cells (GECs), and podocytes (Pod, **Fig. 1i**) and in diabetic eNOS^-/-^ mice, an established murine model with severe DKD (**Supplementary Fig. 2c**,**d**).

In line with the *in vivo* observations, high glucose (HG) concentrations (25 mM) induced p21 expression within 24 hours in human kidney (HEK-293, **Fig. 1j**) and mouse tubular (BUMPT, **Supplementary Fig. 2e**,**f**) cells, and p21 expression remained high despite lowering the glucose concentration to 5 mM for an additional 24 hours in both cell lines. Concomitant exposure of tubular cells (BUMPT) to SGLT2i had no impact on p21 expression (**Supplementary Fig. 2e**,**f**), suggesting that SGLT2i itself does not modulate p21 expression. Congruently, when using insulin to reduce blood glucose levels in the T1DM mouse model, we likewise observed persistent p21 expression in renal cortex extracts and in tubular cells (**Fig. 1k, Supplementary Fig. 2g-j)**.

As p21 is closely associated with senescence (*25*), we performed senescence-associated β-galactosidase (SA-β-gal) staining. SA-β-gal staining was increased in tubular cells of diabetic mice (T1DM and T2DM models) and remained high despite reducing blood glucose levels using SGLT2i or insulin (**Fig. 1l, Supplementary Fig. 3a-c**). Likewise, in renal cells (HEK-293), the maintenance of 25 mM glucose for 24 h induced β-galactosidase staining, which remained high despite returning cells to normal (5 mM) glucose concentrations (**Supplementary Fig. 3d**,**e**). Thus, p21 expression remains elevated despite the normalization of glucose levels and is associated with senescence markers in renal tubular cells in various models of DKD.

### Sustained tubular p21 expression and senescence despite glucose normalization in humans

To scrutinize the relevance of these findings in the context of human DKD, we first analyzed p21 expression by immunohistochemistry. Tubular p21 expression was increased in renal biopsies obtained from diabetic patients with DKD (DM+DKD) but not in diabetic patients without DKD (DM-DKD) or non-diabetic controls (C, **Supplementary Table 1, Fig. 2a**,**b**). Tubular p21 expression was associated with increased γH2A.X staining (phosphorylated H2A histone family member X, a marker of DNA damage and senescence (*26*)) in patients with DKD (**Fig. 2a**,**c**).

**Figure 2:**
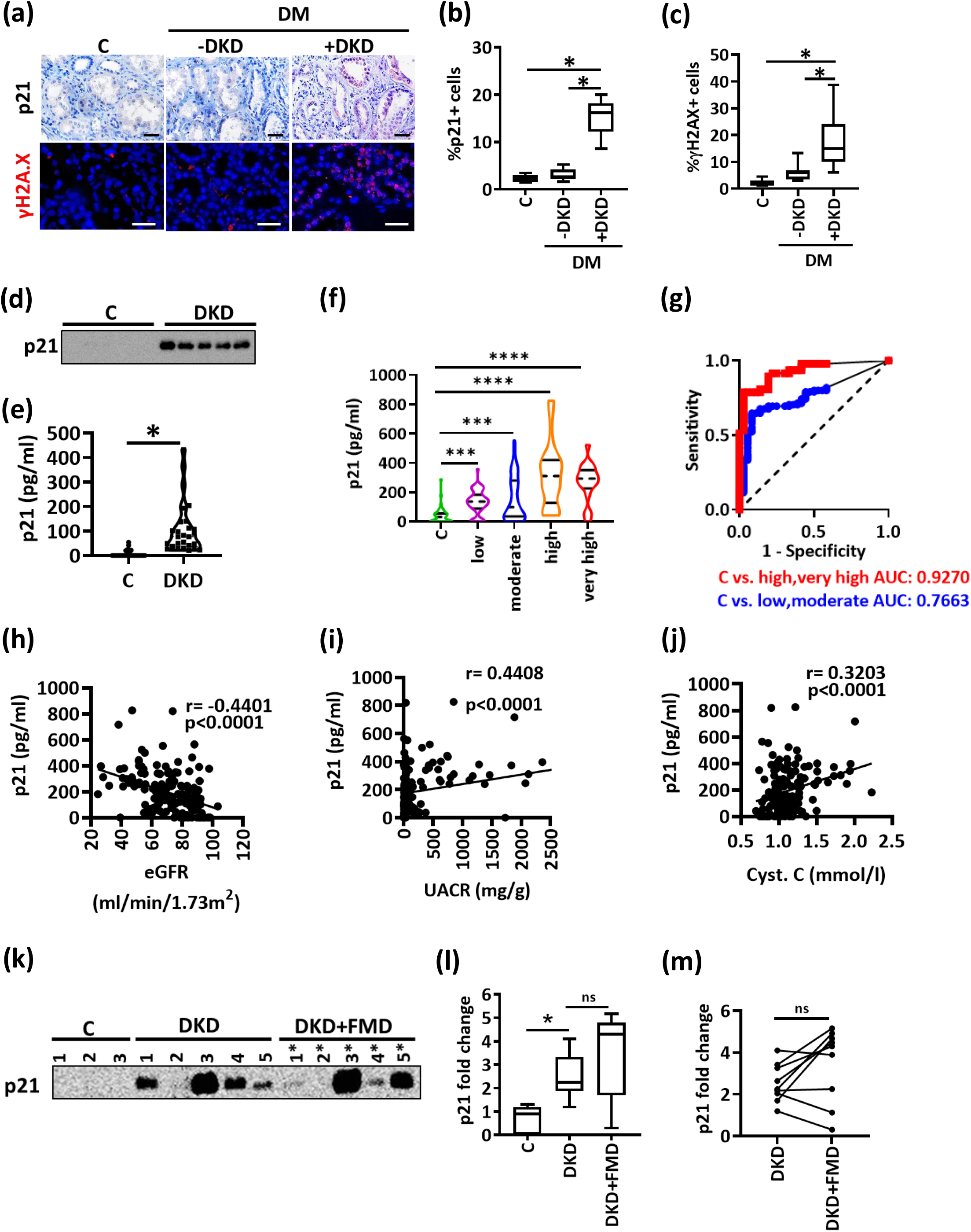
Sustained tubular p21 expression in human DKD. **a-c)** Exemplary histological images of human kidney sections stained for p21 (top) or γH2A.X (histone H2A family X, bottom) obtained from non-diabetic controls (C) or diabetic patients without (DM-DKD) or with (DM+DKD) DKD; p21 is detected by HRP-DAB reaction, brown; hematoxylin nuclear counter stain, blue; γH2A.X is immunofluorescently detected, red; DAPI nuclear counterstain, blue. Box plots summarizing results for p21 (b) and γH2A.X (c); Scale bars represent 20 µm. Box plots reflecting mean±SEM; ANOVA, \**P*<0.05 **d**,**e)** Urinary p21, detected by immunoblotting (d), or ELISA (e; pg/ml) is markedly increased in patients with DKD compared to non-diabetic controls with normal kidney function (C). Exemplary immunoblot (d) and violin plots showing the distribution of the urinary levels of p21 (e). t-test (e) comparing C versus DKD, \**P*<0.05 **f)** Violin plots showing the distribution of the urinary levels of p21 (pg/ml; ELISA) in normoglycemic controls (c) and diabetic individuals which are classified according to KDIGO criteria (risk of CKD: low; moderate; high and very high) from the LIFE-adult cohort. Kruskal-Wallis test, \*\*\**P*<0.001; \*\*\*\**P*<0.0001 **g)** Receiver operating characteristic (ROC) analyses of urinary p21 (pg/ml; ELISA) in diabetic individuals with low or moderate risk of CKD (blue) and in diabetic patients with high or very high risk of CKD (red) compared to non-diabetic controls (C). AUC: area under the curve **h-j)** Negative correlation of urinary p21 (pg/ml, ELISA) with estimated glomerular filtration rate (eGFR (ml/min/1.73m^2^, h) and positive correlation with urinary albumin creatinine ration (UACR, mg albumin /g creatinine, i) or with cystatin C serum levels (Cyt. C, mmol/l, j) in diabetic individuals from the LIFE-adult cohort **k-m)** Exemplary immunoblot (k) of urinary p21 in diabetic patients with known DKD (microalbuminuria) before (DKD) and after fasting mimicking diet (DKD+FMD). Box plot (l) summarizing results and line-graph (m) illustrating individual changes of urinary p21 levels before and after FMD. Box plots (l) reflecting mean±SEM, ANOVA (l) or pearson’s correlation (m), \**P*<0.05; ns: non-significant.

Biopsies from diabetic patients are not routinely obtained, precluding monitoring of tubular p21 expression. Hence, we determined whether p21 can be detected in urine samples. p21 was readily detectable by immunoblotting in randomly chosen urinary samples from diabetic patients with known DKD (N=5), but not in healthy controls (N=5, **Fig. 2d**). We next determined urinary p21 in a group of patients with known DKD (N=26) and compared these to patients with normal renal function (N=22, **Supplementary Table 2**), both recruited from the local outpatient clinic. Urinary p21 was readily detectable by ELISA and on average much higher in patients with DKD (26 out of 26 positive) in comparison to outpatients with other diseases, but normal renal function (6 out of 22 positive, **Fig. 2e**).

To validate this approach, we identified non-diabetic and diabetic individuals in a large cross-sectional cohort (LIFE-Adult, **Supplementary Table 3**) (*27*). The severity of DKD was classified according to the KDIGO criteria (*28*). Urinary p21 was again significantly increased in diabetic participants compared to non-diabetic individuals (**Fig. 2f**). Urinary p21 levels were already increased in diabetic patients with a low or moderate risk of chronic kidney disease (CKD), and further increased in patients with high or very high risk of CKD (**Fig. 2f**). ROC-analyses of urinary p21 revealed an AUC of 0.7663 for the detection of patients at a low or moderate CKD risk, while the AUC was 0.9270 for the detection of patients at a high or very high risk of CKD (**Fig. 2g**), corroborating that urinary p21 increases with the severity of kidney disease. Accordingly, urinary p21 was inversely correlated with the glomerular filtration rate (GFR) and directly correlated with albuminuria and cystatin C serum level in diabetic patients (**Fig. 2h-j**).

To analyze whether p21 levels remain elevated in patients with DKD despite blood glucose reduction, we assessed urinary p21 in diabetic patients before and after blood glucose lowering resulting from a fasting-mimicking diet (**Supplementary Table 4**) (*29*). Urinary p21 was increased in patients with DKD and remained high despite blood glucose reduction in most of these patients (**Fig. 2k-m**).

Taken together, (i) the increase of tubular p21 expression in diabetic patients with DKD, but not in diabetic patients without DKD, (ii) the increase of urinary p21 in diabetic patients in association with an increasing KDIGO grade of chronic kidney disease, and (iii) the persistent increase of urinary p21 despite improved blood glucose levels in diabetic patients support a model in which renal tubular p21 expression reflects persistent tubular damage in DKD, contributing to the hyperglycemic memory. These data raise the question of whether reversing p21 induction may “erase” the hyperglycemic memory.

### aPC reverses glucose induced p21 promoter methylation and sustained p21 expression

We have previously shown that the coagulation protease aPC reverses epigenetically induced p66^Shc^ expression in diabetic vascular diseases (*9, 11*). To determine whether aPC reverses glucose-induced and sustained p21 expression, we conducted *in vitro* experiments using HEK-293 and mouse PTCs. HG induced p21 expression in HEK-293 and PTCs, and the expression remained high despite normalization of glucose levels (HG/NG, **Fig. 3a-c**). Exposure of cells to aPC (20 nM) along with high glucose concentrations had no impact on p21 induction or sustained p21 expression during the subsequent 24 hours of low glucose concentrations (HG(aPC)/NG, **Fig. 3a-b**). In contrast, the addition of aPC at the time of glucose normalization reduced p21 expression (HG/NG(aPC), **Fig. 3a-c**).

**Figure 3:**
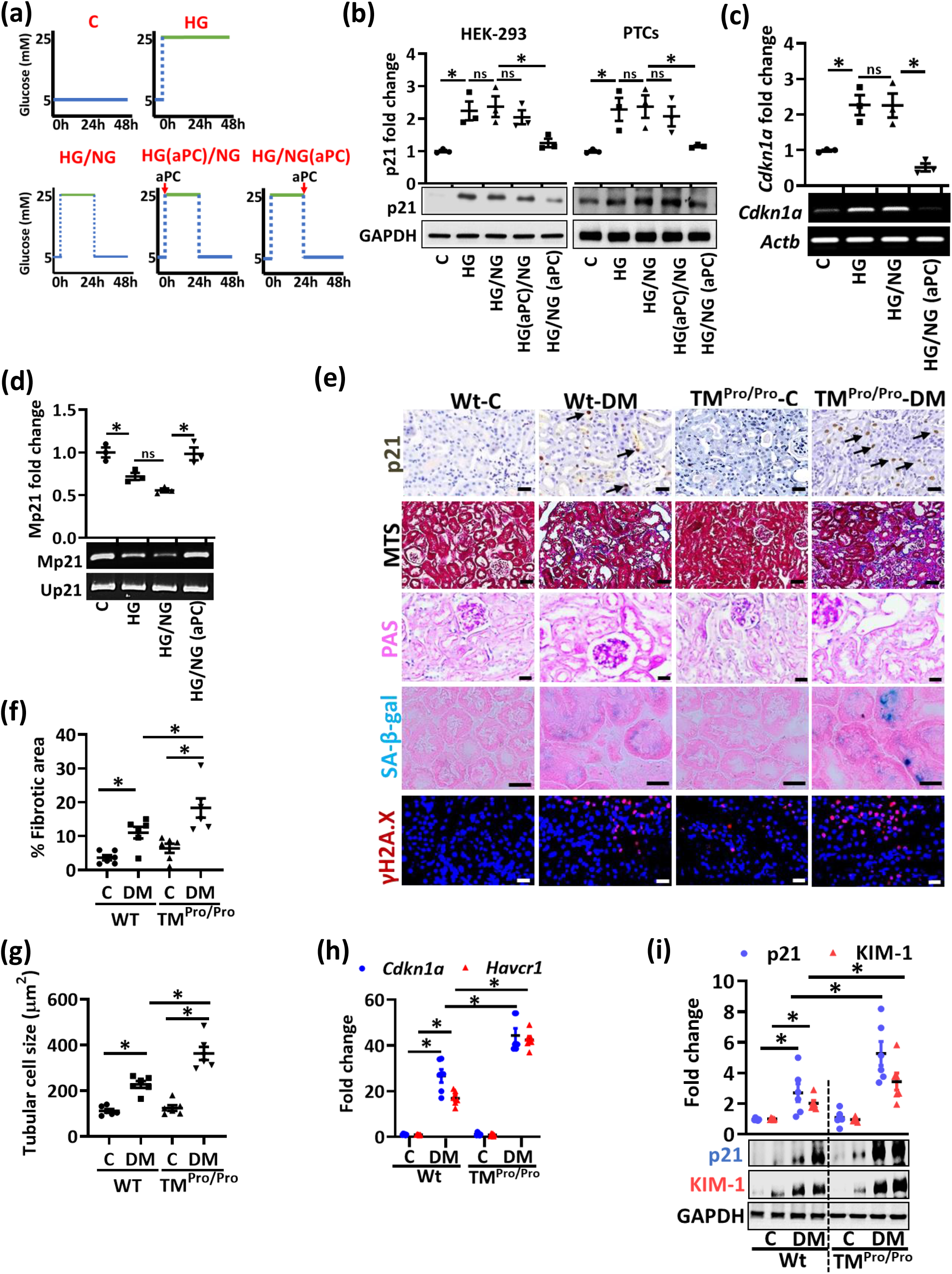
Impaired protein C activation increases tubular p21 expression and senescence *in vivo*. **a)** Experimental scheme of *in vitro* experiments with human kidney cells (HEK-293) or mouse primary tubular cells (PTCs). Experimental conditions: control with continuously normal glucose (C, 5 mM glucose), continuously high glucose (HG, 25 mM, 48 h), high glucose for 24 h followed by normal glucose (NG, 5 mM glucose) for 24 h without aPC-exposure (HG/NG), with aPC (20 nM) during the high glucose exposure (HG(aPC)/NG), or with aPC (20 nM) upon returning cells to normal glucose (HG/NG(aPC)) **b)** Exemplary immunoblots of p21 (bottom; loading control: GAPDH) in HEK-293 cells and mouse PTCs and dot plots summarizing results (top). Experimental conditions as described in a. Dot plots reflecting mean±SEM of at least 3 independent experiments; ANOVA, \**P*<0.05; ns: non-significant **c**,**d)** Exemplary images (bottom) and dot plots summarizing results (top) of p21 (*Cdkn1a*) mRNA-levels determined by semiquantitative RT-PCR (control: β-actin, *Actb*; c) and methylation-specific PCR (d) of the p21-promoter (Mp21; control: unmethylated p21, Up21) in HEK-293 cells(experimental conditions as described in a). Dot plots reflecting mean±SEM of at least 3 independent experiments; ANOVA, \**P*<0.05; ns: non-significant **e)** Exemplary histological images from non-diabetic (C) or diabetic (DM) wild-type (Wt) and TM^Pro/Pro^ mice for p21 (immunohistochemically detected by HRP-DAB-reaction, brown, examples illustrated by arrows; hematoxylin counterstain), interstitial fibrosis (Masson’s trichrome stain, MTS), periodic acid Schiff stain (PAS), SA-β-gal. stain (senescence associated β-galactosidase, blue; eosin counterstain), and γH2A.X immunohistochemistry (histone H2A family X, red; DAPI nuclear counterstain). Scale bars represent 20 μm **f**,**g)** Dot plots summarizing tubular fibrotic area (f) and tubular cell size (g) inexperimental groups (as described in e). Dot plots (f,g) reflecting mean±SEM of at least 6 mice per group; ANOVA, \**P*<0.05 **h)** Dot plot summarizing renal p21 (*Cdkn1a*) and KIM-1 (*Havcr1*) expression (mRNA, qRT-PCR) in experimental groups (as described in e). Dot plot reflecting mean±SEM of at least 6 mice per group; ANOVA, \**P*<0.05 **i)** Representative immunoblots (bottom; loading control: GAPDH) and dot plot summarizing results (top) for renal p21 and KIM-1 expression in experimental groups (as described in e). Dot plot reflecting mean±SEM of at least 6 mice per group; ANOVA, \**P*<0.05.

In view of perpetuated p21 mRNA expression in tubular cells, we next analyzed p21 promoter methylation. HG-induced p21 mRNA expression was associated with reduced p21 promoter methylation (**Fig. 3c**,**d**). Restoring low glucose concentrations had no impact on p21 mRNA expression and p21 promoter hypomethylation (HG/NG, **Fig. 3c**,**d**), while the addition of aPC (20 nM) at the time glucose levels were reduced fully restored p21 promoter methylation and reduced p21 mRNA expression (HG/NG(aPC), **Fig. 3c**,**d**). These results show that aPC reverses glucose-induced, sustained p21 expression and associated p21 promoter hypomethylation in renal tubular cells.

### Impaired protein C activation increases tubular p21 expression and senescence *in vivo*

Hyperglycemia is associated with endothelial dysfunction and impaired thrombomodulin (TM)-dependent protein C activation (*12, 13*). To determine whether impaired TM-dependent protein C activation aggravates glucose-induced tubular p21 expression and senescence, we analyzed mice with low endogenous aPC generation secondary to a point mutation in the TM gene (*Thbd*, Glu404Pro, TM^Pro/Pro^ mice) (*30*). This point mutation mimics impaired TM-dependent protein C activation due to oxidative damage (*31*). If diabetic, TM^Pro/Pro^ mice display increased glomerular damage(*13*), but the consequences for tubular damage and tubular senescence have not been investigated hitherto. In addition to albuminuria and normalized kidney weight, interstitial fibrosis and tubular cell size (reflecting cellular hypertrophy) were enhanced in diabetic TM^Pro/Pro^ mice after 22-weeks of persistent hyperglycemia as compared to diabetic wild-type littermates despite comparable blood glucose levels (**Fig. 3e-g, Supplementary Fig. 4a-e**). Concomitantly, p21 (*Cdkn1a*) and KIM-1 (kidney injury molecule-1, *Havrc1*) expression (mRNA and protein level) were increased in diabetic TM^Pro/Pro^ mice as compared to diabetic wild-type mice (**Fig. 3e**,**h**,**i, Supplementary Fig. 4f**). The increased tubular cell hypertrophy and p21 expression indicate enhanced tubular cell senescence, which was confirmed by greater SA-β-gal and γH2A.X staining in diabetic TM^Pro/Pro^ mice (**Fig. 3e, Supplementary Fig. 4g**,**h**). These data show that impaired endogenous PC activation aggravates tubulointerstitial damage and tubular cell senescence in DKD, potentially in a p21-dependent fashion.

### p21 mediates enhanced tubular senescence in aPC-deficient mice

To analyze the relevance of p21 for increased tubulointerstitial damage and tubular senescence in diabetic aPC-deficient mice, we analyzed TM^Pro/Pro^ mice with genetically superimposed p21 deficiency (TM^Pro/Pro^ x p21^-/-^). Despite comparable blood glucose levels among diabetic mice (22 weeks of persistent hyperglycemia), genetically superimposed p21 deficiency markedly reduced albuminuria and normalized kidney weight in hyperglycemic TM^Pro/Pro^ mice (**Fig. 4a, Supplementary Fig. 5**). In parallel, tubular and interstitial damage (fibrotic area, KIM-1 expression, **Fig. 4b-e**) as well as markers of tubular cell senescence (SA-β-gal and γH2A.Xx, **Fig. 4b**,**f**,**g**) were markedly improved in TM^Pro/Pro^ x p21^-/-^ mice. Remarkably, indices of glomerular injury, such as extracellular matrix accumulation (FMA, fractional mesangial area, **Fig. 4b**,**d**) or reduced nephrin expression (**Fig. 4h**,**i)**, were not improved in TM^Pro/Pro^ x p21^-/-^ mice, indicating that p21 induction causes primarily tubular damage in mice with low aPC levels. Taken together, the increased tubulointerstitial damage and senescence in diabetic TM^Pro/Pro^ mice depends on p21 expression, supporting a model in which thrombomodulin-dependent aPC-generation regulates tubular p21 expression and senescence in DKD.

**Figure 4:**
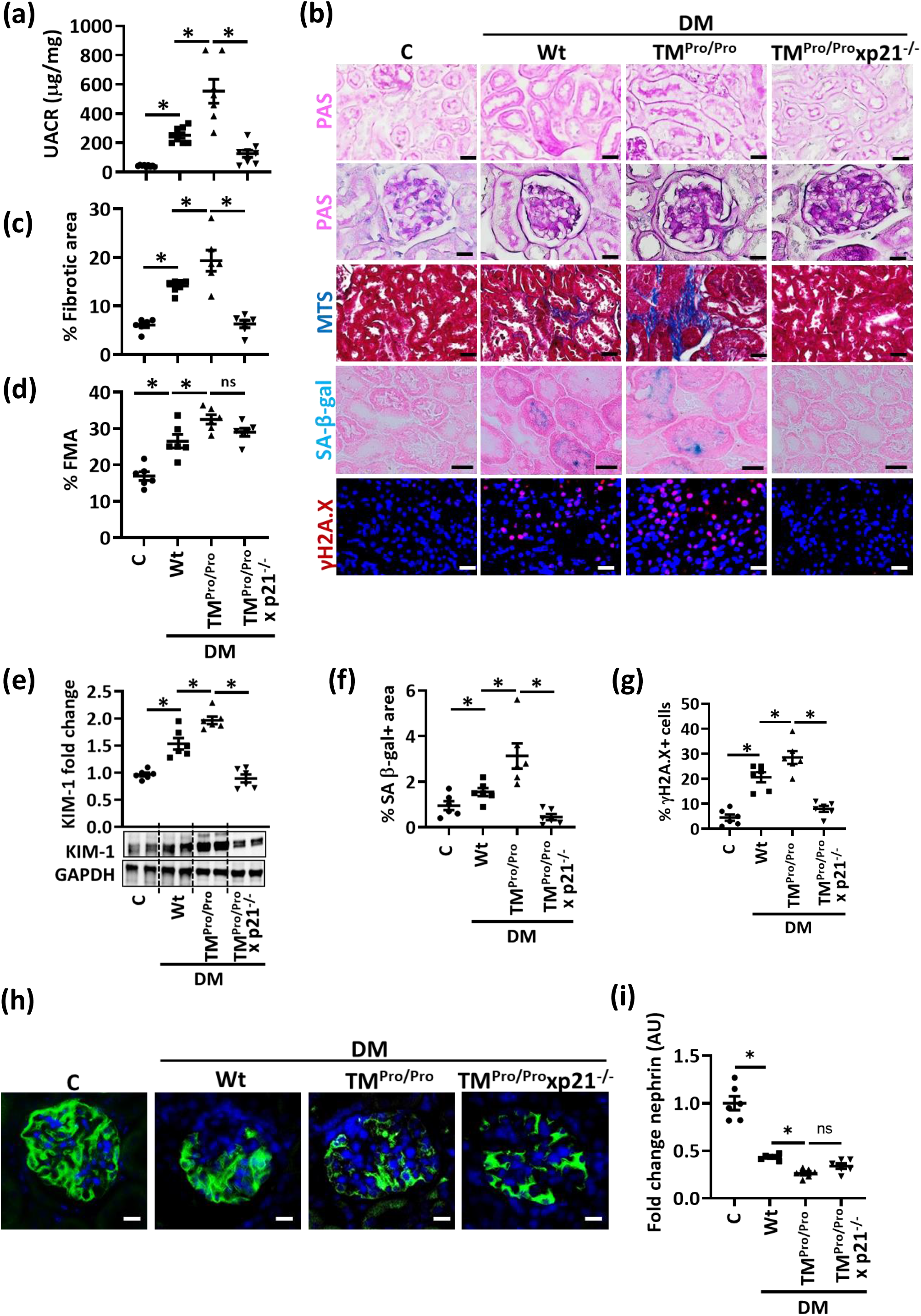
p21 mediates enhanced tubular senescence in aPC-deficient mice. **a)** Dot plot summarizing albuminuria (urinary albumin-creatinine ratio, µg albumin/mg creatinine; UACR) in non-diabetic (C) or diabetic (DM) wild-type (Wt), TM^Pro/Pro^ and TM^Pro/Pro^ x p21^-/-^ mice. Dot plots reflecting mean±SEM of at least 6 mice per group; ANOVA, \**P*<0.05 **b)** Exemplary histological images of periodic acid Schiff stains (PAS) showing tubuli (top) or glomeruli (2^nd^ row), interstitial fibrosis (Masson’s trichrome stain, MTS), SA-β-gal. stain (senescence associated β-galactosidase, blue; eosin counterstain), and γH2A.X immunohistochemistry (histone H2A family X, red; DAPI nuclear counterstain) in experimental groups (as described in a) Scale bars represent 20 μm **c**,**d)** Dot plots summarizing tubular fibrotic area (c) and fractional mesangial area (FMA, d), the latter reflecting glomerulosclerosis in experimental groups (as described in a). Dot plots reflecting mean±SEM of at least 6 mice per group; ANOVA, \**P*<0.05; ns: non-significant **e)** Exemplary immunoblot (bottom; loading control: GAPDH) and dot plot summarizing results (top) of renal KIM-1 expression in experimental groups (as described in a). Dot plots reflecting mean±SEM of at least 6 mice per group; ANOVA, \**P*<0.05 **f**,**g)** Dot plots summarizing percentage of SA-β-gal. positive area (f) and percentage of γH2A.X positive cells (g) in experimental groups (as described in a). Dot plots reflecting mean±SEM of at least 6 mice per group; ANOVA, \**P*<0.05 **h**,**i)** Exemplary histological images of glomerular nephrin expression (green, DAPI nuclear counterstain, blue, h), and dot plot summarizing nephrin staining intensity fold change (i) in experimental groups (as described in a). Dot plot (i) reflecting mean±SEM of at least 6 mice per group; ANOVA, \**P*<0.05; ns: non-significant.

### High glucose and aPC differentially regulate renal DNMT1 expression

We next investigated the mechanism regulating persistent tubular p21 expression and p21-promoter demethylation in hyperglycemic conditions. As DNA methylation depends on the activity of DNA methyltransferases (DNMTs), we analyzed total DNMT activity. HG reduced DNMT activity in HEK-293 cells *in vitro*, which remained low despite normalization of glucose levels (HG/NG), but was restored upon concomitant exposure to aPC (20 nM, HG/NG(aPC), **Fig. 5a**). Expression analyses of individual DNMTs revealed that HG induced sustained suppression of DNMT1 and DNMT3b but had no impact on DNMT3a (**Fig. 5b-d**). Addition of aPC at the time of returning cells to normal glucose concentrations restored DNMT1 and DNMT3b expression *in vitro* (**Fig. 5b**,**d**).

**Figure 5:**
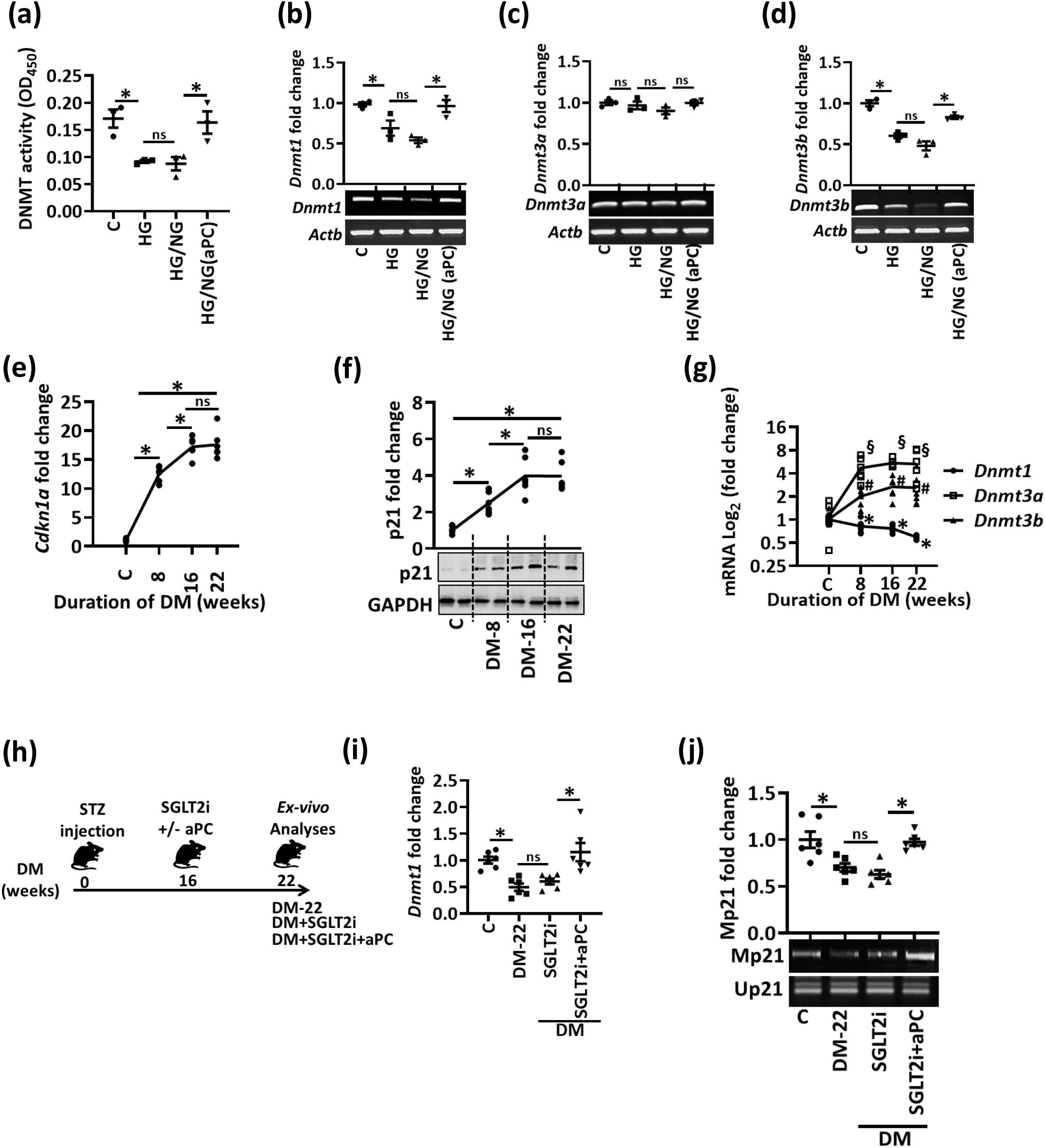
aPC regulates p21 promoter methylation and DNMT1 in tubular cells. **a)** Dot plot summarizing total DNMT-activity in nuclear extracts of human kidney cells (HEK-293). Experimental conditions: control with continuously normal glucose (C, 5 mM glucose), continuously high glucose (HG, 25 mM, 48 h), high glucose for 24 h followed by normal glucose (NG, 5 mM glucose) for 24 h without aPC-exposure (HG/NG), or with aPC (20 nM) upon returning cells to normal glucose (HG/NG(aPC)). Dot plot reflecting mean±SEM of at least 3 independent experiments; ANOVA, \**P*<0.05; ns: non-significant **b-d)** Exemplary gel-images of the expression of *Dnmt1* (b), *Dnmt3a* (c), and *Dnmt3b* (d); (bottom; semiquantitative RT-PCR, loading control: β-actin, *Actb*) and dot plots summarizing results (top) in experimental conditions (as described in a). Dot plots reflecting mean±SEM of at least 3 independent experiments; ANOVA, \**P*<0.05; ns: non-significant **e-g)** Kinetics of renal p21 (*Cdkn1a*) expression (e, mRNA, qRT-PCR; f, protein expression). Exemplary immunoblot (f, bottom; GAPDH: loading control) and line graph summarizing results (top). Line graph summerizing *Dnmt1, Dnmt3a* and *Dnmt3b* expression *in vivo* (g, mRNA, qRT-PCR). Diabetic wild-type mice (STZ model, age 8, 16, or 22 weeks) were compared to non-diabetic mice (C, age 30 weeks). Line graphs reflecting mean±SEM of at least 4 mice per group; ANOVA, * (for *Dnmt1*), # (*Dnmt3b*) and § (*Dnmt3a*) *P*<0.05; ns: non-significant **h)** Experimental scheme: 16 weeks after induction of persistent hyperglycemia with STZ, diabetic mice were treated with PBS (DM-22), sodium/glucose cotransporter 2-inhibitor (Dapagliflozin®, DM+SGLT2i) or a combination of SGLT2i and aPC (DM+SGLT2i+aPC) for further 6 weeks **i**,**j)** Dot plots summarizing renal *Dnmt1* expression (i, mRNA, qRT-PCR), and exemplary images of methylation-specific PCR (j, bottom) and dot plot summarizing results (j, top) of p21-promoter methylation (Mp21; control: unmethylated p21, Up21) in experimental groups (as described in h). Dot plots (i,j) reflecting mean±SEM of at least 6 mice per group. ANOVA; \**P*<0.05; ns: non-significant.

To scrutinize the relevance of p21 and DNMTs *in vivo*, we first determined the expression of p21 and DNMTs at different time points after the induction of persistent hyperglycemia. An increase of p21 mRNA and protein expression was already detectable 8 weeks after induction of hyperglycemia and increased further until week 16 (**Fig. 5e**,**f**). In parallel, we observed a reduction in DNMT1 *in vivo*, whereas DNMT3a and DNTM3b were induced (**Fig. 5g**). These data support a role of DNMT1 in regulating p21 expression in DKD.

To determine whether DNMT1 suppression is memorized in diabetic mice and whether aPC can reverse memorized DNMT1 suppression *in vivo*, we again reversed hyperglycemia in mice after 16 weeks for an additional 6 weeks (DM+SGLT2i, **Fig. 5h**). A subgroup of mice received aPC in addition to the SGLT2i following an established protocol (DM+SGLT2i+aPC, **Fig. 5h**) (*9*). In mice with persistent hyperglycemia for 22 weeks (DM-22), DNMT1 expression was reduced (**Fig. 5i**). A reduction of blood glucose levels by SGLT2i from weeks 16 to 22 did not restore DNMT1 expression (DM+SGLT2i, **Fig. 5i**), while addition of aPC restored DNMT1 expression (DM+SGLT2i+aPC, **Fig. 5i**). Parallel to restoring DNMT1 expression, aPC reversed hyperglycemia-induced persistent hypomethylation of the p21 promoter (**Fig. 5j**). These results support a model in which reduced DNMT1 expression is linked with p21 promoter hypomethylation and increased p21 expression in DKD, which can be reversed by aPC.

### aPC reverses glucose-induced and sustained renal p21 expression and tubular injury *via* DNMT1

To analyze whether aPC reverses tubular injury and senescence *via* DNMT1 *in vivo*, we again reduced blood glucose levels in diabetic mice using SGLT2i starting at week 16 post induction of hyperglycemia for further 6 weeks (DM+SGLT2i), adding aPC in a subgroup of mice (DM+SGLT2i+aPC+PBS, **Fig. 6a**,**b**). In agreement with results shown in Fig.1, lowering blood glucose levels did not normalize albuminuria (**Fig. 6c**). Likewise, markers of tubular injury (tubular dilatation, tubulointerstitial fibrosis, increased KIM-1 expression) and tubular senescence (p21 expression, SA-β-gal and γH2A.X) as well as reduced DNMT1 expression were comparable in SGLT2i-treated (DM+SGLT2i) mice and mice with persistent hyperglycemia (DM-22, **Fig. 6d-f, Supplementary Fig. 6**). In contrast, treatment of diabetic mice with aPC in addition to SGLT2i from weeks 16 to 22 markedly reduced albuminuria to levels below those observed after 16 weeks of hyperglycemia (DM-16 versus DM+SGLT2i+aPC+PBS, **Fig. 6b**) and comparable to those in non-diabetic control mice (C versus DM+SGLT2i+aPC+PBS, **Fig. 6b**). In addition, markers of tubular damage (tubular dilatation, tubulointerstitial fibrosis, KIM-1 expression) and tubular senescence (SA-β-gal, γH2A.X, and p21 expression) were markedly reduced while DNMT1 expression was increased by aPC (**Fig. 6d-f, Supplementary Fig. 6**).

**Figure 6:**
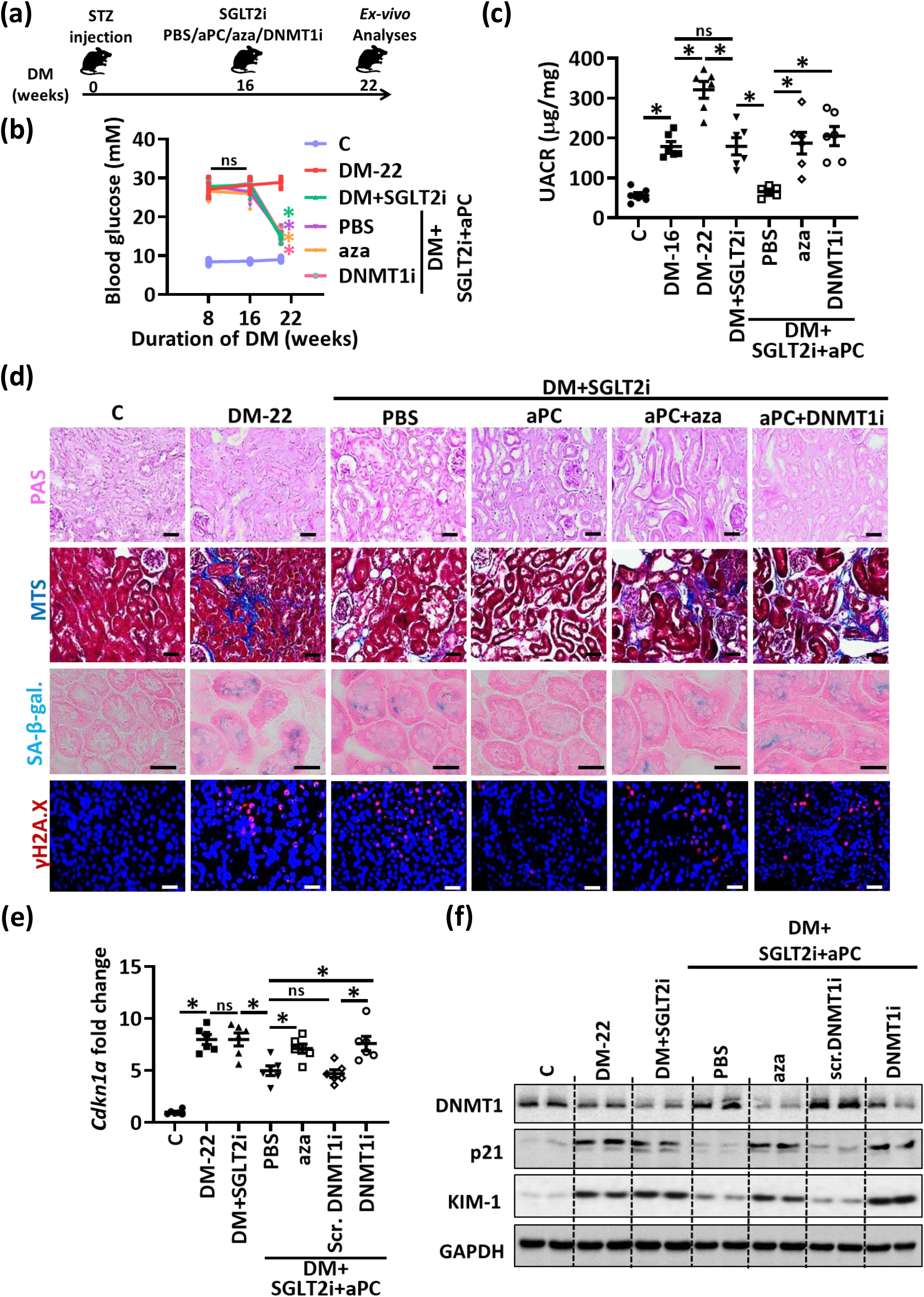
aPC reverses epigenetically sustained renal p21 expression and tubular senescence. **a)** Experimental scheme: 16 weeks after induction of persistent hyperglycemia, diabetic mice were treated with PBS (DM-22), sodium/glucose cotransporter 2-inhibitor (Dapagliflozin®, DM+SGLT2i), a combination of SGLT2i and aPC (DM+SGLT2i+aPC), a combination of SGLT2i, aPC and the pan DNMTs-inhibitor 5-aza-deoxycytidine (DM+SGLT2i+aPC+aza), or a combination of SGLT2i, aPC and a vivo morpholino targeting DMNT1 (DM+SGLT2i-aPC-DNMT1i) for further 6 weeks **b)** Average blood glucose levels in experimental groups (as described in a) after 8 or 16 weeks of persistent hyperglycemia and at 22 weeks. Line graphs reflecting mean±SEM of at least 6 mice per group; ANOVA, \**P*<0.05; ns: non-significant **c)** Dot plot summarizing albuminuria (urinary albumin-creatinine ratio, µg albumin/mg creatinine; UACR) in experimental groups (as described in a). Baseline albuminuria before interventions was determined after 16 weeks of hyperglycemia (DM-16). Dot plot reflecting mean±SEM of at least 6 mice per group. ANOVA; \**P*<0.05; ns: non-significant **d)** Exemplary histological images of periodic acid Schiff stains (PAS), interstitial fibrosis (Masson’s trichrome stain, MTS), SA-β-gal. stain (senescence associated β-galactosidase, blue; eosin counterstain), and γH2A.X (histone H2A, family X, immunohistochemistry, red; DAPI: nuclear counterstain) in experimental groups (as described in a); scale bars represent 20 μm **e**,**f)** Expression of renal p21 mRNA (*Cdkn1a*, qRT-PCR, e) and p21, DNMT1 and KIM-1 protein (exemplary immunoblot, f; loading control: GAPDH in experimental groups (as described in a). A nonspecific scrambled vivo morpholino (scr. DNMT1i) has no effect. Dot plot (e) reflecting mean±SEM of at least 6 mice per group. ANOVA; \**P*<0.05; ns: non-significant.

To determine whether aPC controls p21 expression and tubular injury via DNMT1, we chose two complimentary approaches. We either (i) added the pan DNMT inhibitor 5-aza-2’-deoxycytidine (DM+SGLT2i+aPC+aza; 5-aza-2’-deoxycytidine reduces DNMT1 activity and its expression (*32*)) or (ii) specifically inhibited DNMT1 expression (using vivo-morpholino (*33*), DM+SGLT2i+aPC+DNMTi) on top of SGLT2i and aPC treatment in mice with 16 weeks of established hyperglycemia (**Fig. 6a**). 5-aza-2’-deoxycytidine and DNMT1 vivo morpholino markedly reduced DNMT1 expression (**Fig. 6f, Supplementary Fig. 6g)**. Both interventions abolished aPC’s protective effect, increasing albuminuria levels, tubular dilatation, tubulointerstitial fibrosis, KIM-1 and p21 expression and the senescence markers SA-β-gal and γH2A.X to the levels observed in mice treated with SGLT2i only (**Fig. 6c-f, Supplementary Fig. 6**). In agreement with an epigenetic regulation, p21 mRNA levels were suppressed by aPC (DM+SGLT2i+aPC+PBS), but this effect was reversed upon additional treatment with 5-aza-2’-deoxycytidine (DM+SGLT2i+aPC+aza) or DNMT1 vivo morpholino (DM+SGLT2i+aPC+DNMTi, **Fig. 6e**). A nonspecific scrambled vivo morpholino (Scr-DNMTi) had no effect (**Fig. 6e**,**f, Supplementary Fig. 6g** and data not shown). These data establish that aPC controls renal p21 expression *via* DNMT1.

### aPC regulates epigenetically sustained p21 expression independent of its anticoagulant function

aPC conveys its cytoprotective effect at least partially independent of its anticoagulant properties through receptor-dependent mechanisms, involving typically protease-activated receptors (PARs) and the endothelial protein C receptor (EPCR) (*34*). Expression of PAR1, PAR2, PAR4, and EPCR, known to be expressed in an immortalized mouse tubular cell line (BUMPT-cells) (*35*), was confirmed in mouse PTCs (**Supplementary Fig. 7a**). Upon knockdown of PARs or EPCR, we identified PAR1 and EPCR as the receptors required for aPC-mediated reversal of glucose-induced and sustained p21 and KIM-1 expression (**Supplementary Fig. 7b**,**c**). The receptor dependent modulation of p21 expression suggests that aPC’s effect on tubular p21 expression is independent of its anticoagulant function.

Do determine whether aPC reverses glucose induced p21 expression and associated tubular damage independent of its anticoagulant function *in vivo*, we used aPC-based pharmacological agents with markedly reduced anticoagulant potential (*9*): (i) 3K3A-aPC, an aPC mutant with largely reduced anticoagulant but sustained signaling efficacy and (ii) parmodulin-2, a small peptide mimicking biased aPC signaling *via* PAR1 (*36-38*). Again, after 16 weeks of hyperglycemia we initiated treatment with SGLT2i in combination with phosphate-buffered saline (PBS, control), with 3K3A-aPC, or with parmodulin-2 (**Fig. 7a**). While blood glucose levels in SGLT2i plus PBS, 3K3A-aPC or parmodulin-2 treated mice were comparable (**Fig. 7b**), 3K3A-aPC or parmodulin-2 on top of SGLT2i both markedly reduced albuminuria compared to that in SGLT2i+PBS mice (**Fig. 7c**). While SGLT2i+PBS had no impact on DNMT1 or p21 expression, 3K3A-aPC and parmodulin-2 in addition to SGLT2i increased DNMT1 while reducing p21 expression in renal cortex extracts (**Fig. 7d-f, Supplementary Fig. 8a**,**b**), which was associated with improvement of renal tubular damage (reduced tubular dilatation, interstitial fibrosis, KIM-1 expression) and a reduction of senescence markers (tubular SA-β-gal and γH2A.X expression, **Fig. 7f**,**g and Supplementary Figure 8c-f**).

**Figure 7:**
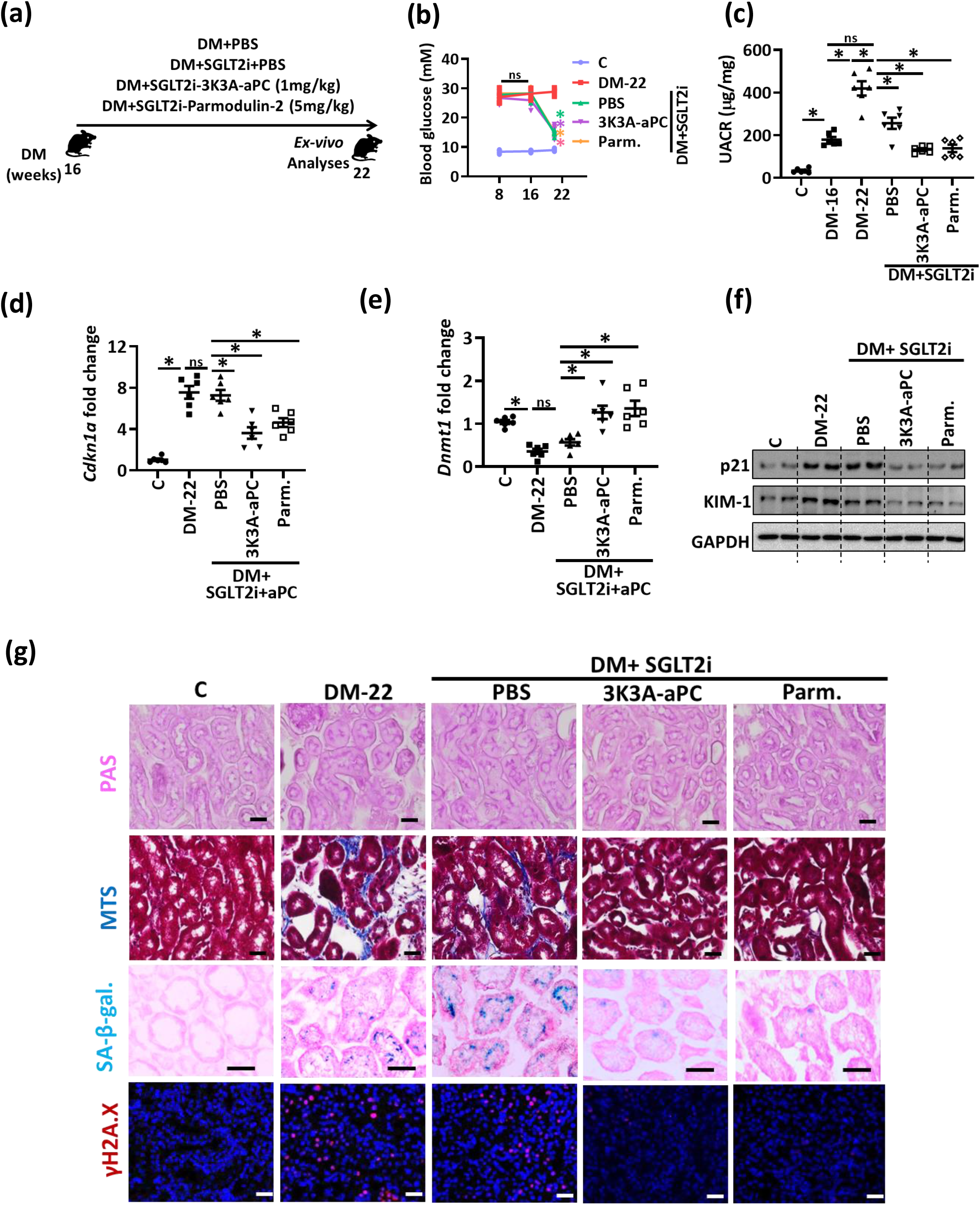
aPC regulates epigenetically sustained p21 expression independent of its anticoagulant function. **a)** Experimental scheme: 16 weeks after induction of persistent hyperglycemia, diabetic mice were treated with PBS (DM+PBS), sodium/glucose cotransporter 2-inhibitor (Dapagliflozin®, DM+SGLT2i), a combination of SGLT2i and 3K3A-aPC (DM+SGLT2i+3K3A-aPC) or a combination of SGLT2i, aPC and parmodulin-2 (DM+SGLT2i+Parm.) for further 6 weeks **b)** Average blood glucose levels after 8 and and16 weeks of persistent hyperglycemia and at 22 weeks in experimental groups (as described in a). Line graphs reflecting mean±SEM of at least 6 mice per group; ANOVA, \**P*<0.05; ns: non-significant **c)** Dot plot summarizing albuminuria (urinary albumin-creatinine ratio, µg albumin/mg creatinine; UACR) in experimental groups (as described in a). Baseline albuminuria before interventions was determined after 16 weeks of hyperglycemia (DM-16). Dot plot reflecting mean±SEM of at least 6 mice per group. ANOVA; \**P*<0.05; ns: non-significant **d-f)** Expression of renal p21 mRNA (*Cdkn1a*, qRT-PCR, d; and protein, f), renal DNMT1 mRNA (*Dnmt1*, qRT-PCR, e), and renal KIM-1 (protein, f) in experimental groups (as described in a). Exemplary immunoblots (f; GAPDH: loading control). Dot plots reflecting mean±SEM of at least 6 mice per group. ANOVA; \**P*<0.05; ns: non-significant **g)** Exemplary histological images of periodic acid Schiff stain (PAS), interstitial fibrosis (Masson’s trichrome stain, MTS), SA-β-gal. stain (senescence associated β-galactosidase), and γH2A.X (histone H2A, family X, red, immunofluorescent, DAPI nuclear counterstain) in experimental groups (as described in a); scale bars represent 20 μm.

Taken together, these data demonstrate that sustained p21 expression drives hyperglycemic memory in the context of DKD. Therapeutic approaches either mimicking aPC signaling reduce tubular p21 expression and reverse – at least in mice – hyperglycemic memory (**Fig. 8**).

**Figure 8:**
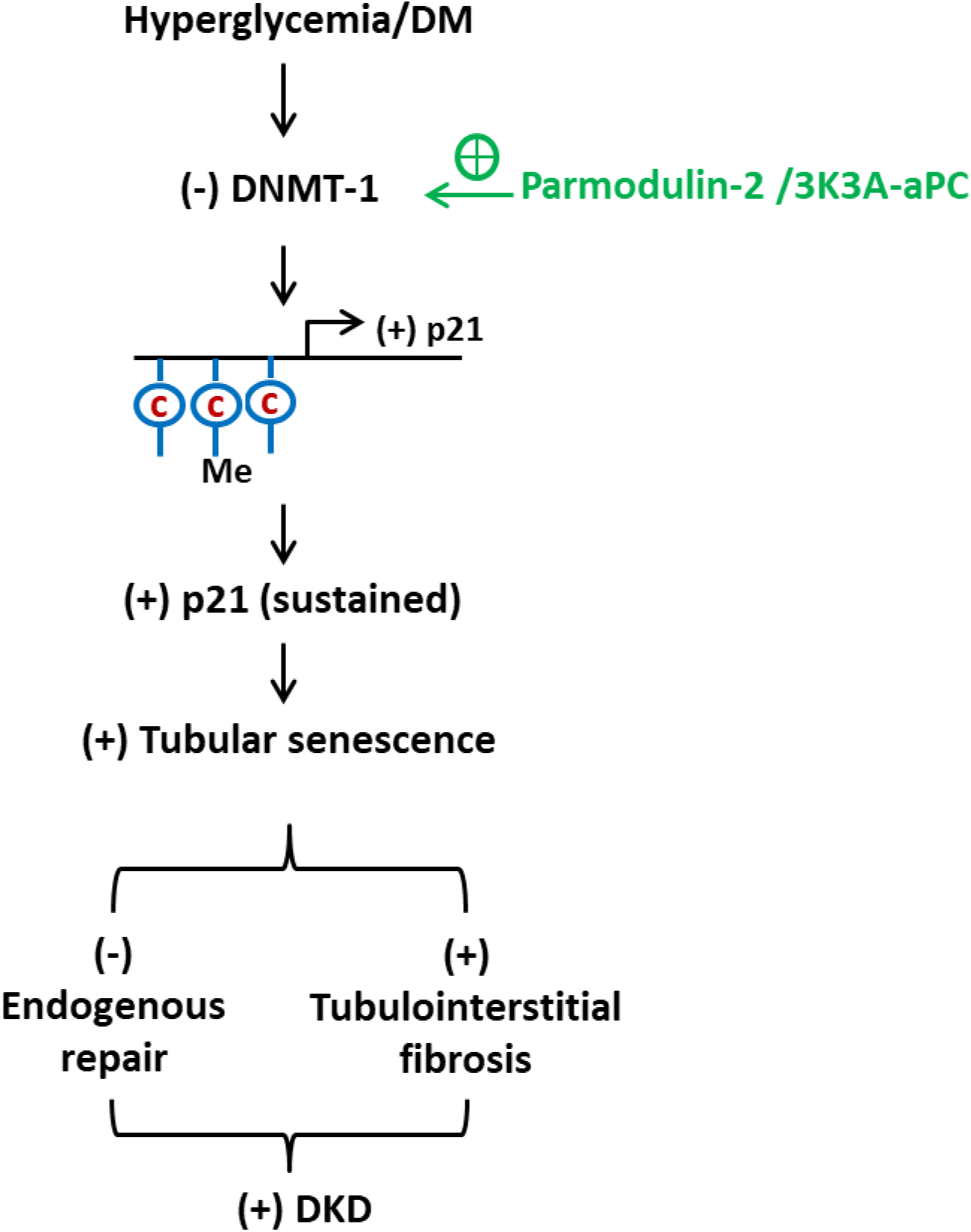
Proposed model summarizing the p21-dependent regulation of the hyperglycemic memory and possible interventions: Hyperglycemia suppresses DNMT1 expression in the tubular compartment. This leads to hypomethylation of the p21 promoter, increased p21 expression, and enhanced tubular senescence. The latter impairs the endogenous tubular repair capability while inducing tubulointerstitial fibrosis. This leads to tubulointerstitial damage and contributes to DKD. Exploiting aPC-signaling (Parmodulin, 3K3A-aPC) reverses the epigenetically sustained p21 expression.

## Discussion

We conducted an unbiased expression analysis aiming to define pathways contributing to hyperglycemic memory in DKD, e.g., the persistence of albuminuria despite a marked improvement in blood glucose levels. Approximately half of the hyperglycemia-induced changes in gene expression persisted despite blood glucose reduction, reflecting hyperglycemic memory. The senescence-associated cyclin-dependent kinase inhibitor p21 (*Cdkn1a*), a gene regulating cell proliferation and senescence (*39*), was among the top genes that were persistently induced. p21 itself is controlled by DNMT1. Reducing p21, either by restoring DNMT1 or exploiting aPC signaling, reverses already established DKD in mice (**Fig. 7**). This study identified p21-dependent tubular senescence as a pathway contributing to hyperglycemic memory and demonstrates that this mechanism can be therapeutically targeted.

p21 has been linked with DKD previously, but the regulation of p21 and its role in hyperglycemic memory are not known (*40-43*). The role of p21 in tubular senescence in DKD is in agreement with previous observations: (i) the level and duration of p21 induction determine the onset of cellular growth arrest, (ii) p21 expression is sufficient to induce senescence, (iii) p21 expression restricted to tubular cells is sufficient to induce renal fibrosis following acute kidney injury (*44-46*), and (iv) p21 induces endoreduplication (*47*), which may impair renal tubular cell recovery following acute kidney injury (*48, 49*). These and the current data support a model in which prolonged and sustained p21 expression causes tubular senescence and compromises tubular repair capacity in DKD patients (*50*), even after glycemic control has improved.

The current results suggest that epigenetic regulation of p21 depends on DNMT1. DNMT1 is a regulator of tubular development and nephron progenitor cell renewal during embryonic nephrogenesis (*51, 52*), supporting a crucial function of DNMT1 in the renal tubular compartment. However, DNMT1 is not expected to specifically target p21 and its reduced expression in DKD is expected to de-repress (hence induce) other genes. Accordingly, about 93% of genes induced in hyperglycemic mice remained elevated despite lowering blood glucose levels, while only about 36% of genes suppressed in hyperglycemic mice remained repressed (**Fig. 1 d**), supporting a concept in which reduced DNMT1 activity results in DNA-demethylation and increased expression of various genes. Analyses of human kidney tissue and peripheral blood monocytes identified hypomethylation and increased expression of other genes previously linked with DKD, such as the integrin β2, genes related to TNF signaling and the redox-regulator TXNIP, corroborating that DNA hypomethylation contributes to the persistent expression of genes linked with diabetic vascular complications (*53, 54*). Hence, we cannot exclude that other genes regulated by DNMT1 contribute to the hyperglycemic memory. Likewise, it is possible that persistent hypermethylation and reduced gene expression contribute to the hyperglycemic memory. Yet, the current data do support a crucial function of p21 for the hyperglycemic memory.

The regulation of DNMT1 by aPC raises the intriguing possibility that this coagulation protease may epigenetically control other genes. Indeed, in addition to its effect on p21, aPC reverses glucose-induced expression of p66^Shc^, a mitochondrial protein promoting ROS-generation, in podocytes (glomerular cells) and macrophages (*9, 11*). Furthermore, inhibitors of coagulation factors fIIa (dabigatran) or fXa (rivaroxaban) differentially regulate aPC generation, which is linked to differential gene expression and epigenetic marks (*55*). Epigenetic control of gene-expression by aPC provides a rational for the long-lasting effects of therapeutic aPC applications despite its short half-life (*56*) and – considering the broad cytoprotective effects of aPC – may be relevant in other disease settings (*34, 57*).

The reversal of glucose-induced and sustained p21 or p66^Shc^ expression in tubular cells and in podocytes, respectively, is in agreement with the engagement of different receptor complexes and different mechanisms regulated by aPC in the tubular and glomerular compartments. Thus, aPC signals *via* a heterodimer of PAR3 with PAR2 (human) or PAR1 (mouse) in conjunction with integrin β_3_ in podocytes, while in tubular cells aPC signals *via* a heterodimer of PAR1 and EPCR (*9, 58-60*) (and current results). Mechanistically, aPC targets the unfolded protein response in glomerular cells (*61*) and cellular senescence in tubular cells (this study). These observations support a concept in which different pathomechanisms in the glomerular and tubular compartments promote DKD (*62, 63*). Importantly, aPC ameliorates both glomerular and tubular damage despite targeting different and cell-specific receptor-complexes and pathways, providing a rationale for its high efficacy in animal models of renal injury (*9, 58-60*) (and current results).

aPC has shown beneficial effects in a number of acute and chronic disease models(*34, 56*), suggesting that this coagulation protease is a disease-resolving mediator. However, the therapeutic value of aPC itself is limited by its inherent anticoagulant properties. As previously shown in acute disease models(*36, 64*) and here for the first time in a chronic disease model, harnessing specifically the signaling-dependent disease-resolving effects of aPC is feasible by mimicking its signaling properties. aPC mutants lacking binding sites for the coagulation factor Va (3K3A-aPC) or small molecules mimicking biased signaling of aPC *via* the G-protein coupled receptor PAR1(*36, 56*) (parmodulins) are devoid of an anticoagulant function, rendering aPC-based therapies feasible not only in acute, but also chronic diseases.

We used established therapeutic approaches to improve hyperglycemia (SGLT2i and insulin) and observed sustained p21 expression in both cases, supporting the notion that sustained p21 expression reflects the hyperglycemic memory independent of the glucose-lowering intervention. Lowering blood glucose levels provided partial protection, as reflected by halted disease progression, which is in agreement with the protective effects of SGLT2 inhibitors in clinical studies (*65*). Intriguingly, aPC-based therapeutics provided a benefit on top of SGLT2-inhibtion. Therefore, despite recent data suggesting that SGLT2 inhibitors are useful new therapeutic adjuncts in patients with diabetes, additional therapies for full kidney recovery may be required.

To circumvent the lack of kidney biopsies from diabetic patients before and after improved blood glucose control, we determined urinary p21 levels. p21 was strongly increased in the urine of most diabetic patients and increased with the severity of DKD. Distribution of p21 levels in advanced stages of DKD appeared to be dichotomic, with a small subgroup of patients presenting with low urinary p21 levels despite kidney dysfunction (Fig. 2e). Considering the proposed role of p21 as a mediator and biomarker of the hyperglycemic memory in DKD, urinary p21 levels may allow better stratification of patients with DKD, identifying those patients with advanced kidney injury and an established renal hyperglycemic memory (*66*). If confirmed, these patients may not respond to glucose lowering alone, but may profit from a potential therapy targeting tubular senescence.

In summary, we identified a pathway contributing to hyperglycemic memory in the kidney and provided experimental *in vivo* evidence that targeting hyperglycemic memory in the context of DKD is feasible, potentially by mimicking the cytoprotective effects of aPC (e.g., using small molecules such as parmodulins, pepducins, or aptamers) (*36, 56, 67*).

## Methods

### Reagents

The following antibodies were used in the current study: rabbit anti-p21 (sc-397; Santa Cruz Biotechnology, Germany); rabbit anti-DNMT1 (5032), rabbit anti-human p21 (2947), rabbit anti-γH2A-X (9718), rabbit anti-β-actin (4970) and goat anti-rabbit IgG HRP-linked (7074) (Cell Signaling Technology, Germany); rabbit anti-KIM-1 (ab47635) and rabbit anti-mouse p21 (ab188224) (abcam, USA); rabbit anti-GAPDH (G9545) (Sigma-Aldrich, Germany); goat anti-mouse nephrin (AF3159; R&D systems, USA). The following secondary antibodies for immunofluorescence were used: texas red goat anti-rabbit IgG (TI-1000) and fluorescein rabbit anti-goat IgG (FI-5000), (Vector Laboratories, USA).

Other reagents: Dapagliflozin® (Santa Cruz Biotechnology, Germany); Dulbecco′s Modified Eagle′s Medium (DMEM), DMEM/F-12, trypsin-EDTA, penicillin/streptomycin solution, fetal bovine serum (FBS), ITS supplement, D(+)-Glucose, type I collagenase, RNA*later*™, Hank’s Balanced Salt Solution (HBSS) and HEPES buffer (PAA Laboratories, Austria); donkey serum, horse serum, antibiotic/antimycotic solution, APO transferrin and hydrocortisone solution (Sigma-Aldrich, Germany); recombinant human epidermal growth factor (R&D systems, USA); IFN-γ (Cell Sciences, Germany); BCA reagent (Perbio Science, Germany); vectashield mounting medium with DAPI and citrate-based antigen-unmasking solution (citrate-based, Vector Laboratories, USA); PVDF membrane and immobilon TM western chemiluminescent HRP substrate (Millipore, United States); ammonium persulphate (Merck, Germany); streptozotocin (Enzo Life Sciences, Germany); insulin glargine (Lantus®, Sanofi, France); accu-chek test strips, accu-check glucometer and protease inhibitor cocktail (Roche Diagnostics, Germany); albumin fraction V, hematoxylin Gill II, acrylamide, agarose, periodic acid-Schiff (PAS) reagent, aqueous Eosin, Histokitt synthetic mounting medium and dimethylsulfoxide (DMSO, Carl ROTH, Germany); aqueous mounting medium (ZYTOMED, Germany); Trizol Reagent, soyabean trypsin inhibitor and PBS (Life Technologies, Germany); mouse DNMT1 vivo morpholino and DNMT1 mismatch vivo morpholino (Gene Tools, USA).

Kits used in this study: RevertAid™ H minus first strand cDNA synthesis kit, Dynabeads® mRNA DIRECT™ micro purification kit (Thermo Fisher Scientific, Germany); QIAamp DNA mini kit, RNeasy mini kit, EpiTect MSP kit; EZ DNA Methylation kit (Zymo Research, Germany), human p21 matched antibody pair kit (abcam, USA); mouse albumin quantification ELISA kit (Bethyl Laboratories, Germany); Trichrome stain (Masson) kit and senescence cells histochemical staining kit (Sigma-Aldrich, Germany); DAB substrate kit (Vector Laboratories, USA); EpiQuik DNA methyltransferase activity/inhibition assay kit (Epigentek, USA); ScriptSeq v2 RNA-Seq library preparation kit (Epicentre Technologies, USA), TruSeq SBS Kit v3-HS (Illumina, USA).

### Mice

Wild-type C57BL/6 and p21^*CIP1/WAF1*^ constitutive knock-out (p21^-/-^) mice were obtained from Jackson Laboratory (Bar Harbor, ME, USA). Non diabetic C57BLKsJ-db/+ (db/m) and diabetic C57BL/KSJ-db (db/db) mice were obtained from Janvier (S.A.S., St. Berthevin Cedex, France). Age and sex-matched mice were used in the study. TM^Pro/Pro^ mice have been previously described (*13, 30*). TM^Pro/Pro^ mice were crossed with p21^-/-^ mice, to generate TM^Pro/Pro^ x p21^-/-^ double mutant mice. Mice had been backcrossed onto the C57BL/6 background for at least 9 generations and were routinely maintained on the C57BL/6 background. Age matched littermates were randomly assigned to intervention or control groups.

In some mice hyperglycemia was induced using low dose streptozotocin (STZ) injection following established protocols (intraperitoneally, 60 mg/kg body weight, freshly dissolved in 0.05 mol/l sterile sodium citrate, pH 4.5 for 5 consecutive days) (*9, 58, 68, 69*). Age-matched control mice received sodium-citrated intraperitoneally for five consecutive days. Blood glucose and body weight were measured once weekly (*9, 13*). On average 85 - 90% of STZ-injected mice became diabetic (blood glucose >17 mmol/l) within the first 4 weeks and these were included as diabetic mice in the experiments. Mice that did not develop persistently elevated blood glucose levels and maintained blood glucose levels <11 mmol/l despite STZ-injections were included in the control group (*13*). The endpoints analysed did not differ between STZ-injected, but normoglycaemic mice and sodium-citrated injected mice (controls). Blood and tissue samples were collected after 22 weeks of persistent hyperglycemia (*9, 13, 58, 68, 69*). All animal experiments were conducted following standards and procedures approved by the local Animal Care and Use Committee (Landesverwaltungsamt Halle and Leipzig, Germany).

### *In vivo* intervention studies

Subsets of mice were treated with a sodium-glucose co-transporter 2 inhibitor (SGLT-2i, Dapagliflozin®, 25 mg/kg body weight, in the drinking water (*69*), with the activated protein C (aPC, 1 mg/kg body weight, every other day, intraperitoneally) (*9, 61, 70*), with insulin glargine (Lantus®, 1-2 IU, every other day, subcutaneously), with 5-aza-2’-deoxycytidine (aza, 0,1 mg/kg body weight, every other day, intraperitoneally) (*11, 71*), an aPC variant lacking specifically anticoagulant function (3K3A-aPC; 1 mg/kg body weight, every other day, intraperitoneally) (*36*), the biased PAR1 agonist parmodulin-2 (ML161, Parm, 5 mg/kg body weight, every other day, intraperitoneally) (*36, 38*), DNMT1 vivo morpholino (DNMTi, 5’-CAG GTT GCA GAC GAC AGA ACA GCT C-3’, targeting the translation of transcript variant 1 of mouse DNMT1; 6 mg/kg body weight in PBS, every other day, intraperitoneally) (*72*), or DNMT1 mismatch vivo morpholino (Scr.DNMT1i, 5’-CAG CTT CCA GAC CAC ACA ACA CCT C-3’, 6 mg/kg body weight in PBS, every other day, intraperitoneally). Control mice received PBS every other day (intraperitoneally). All interventions were conducted for 6 weeks, starting after 16 weeks of persistent hyperglycemia. Albuminuria indicative of DKD was ensured before initiating treatment.

### Cell culture and *in vitro* interventions

Human embryonic kidney cells (HEK-293) were routinely grown and maintained in DMEM low glucose (5.5 mM glucose concentration) medium in the presence of 10% FBS at 37°C according to manufacturer’s instructions (ATCC, USA). Mouse primary proximal tubular cells (PTC) were isolated and cultured in DMEM/F-12 medium according to an established protocol (*35, 73*). Immortalized Boston University mouse proximal tubular cells (BUMPT) were cultured in DMEM low glucose in the presence of 10% FBS and maintained at 33°C in the presence of interferon γ (INF-γ, 10 U/ml) to enhance the expression of a thermosensitive T-antigen (*35, 74*). Under these conditions BUMPT cells remain proliferative. To induce the differentiation, BUMPT cells were grown for 2 days at 37°C in the same medium without INF-γ. Differentiation was ascertained by expression of megalin, a marker of proximal tubular cells. Only differentiated cells were used for further experiments.

The *in vitro* model of hyperglycemic memory was performed as described before (*75*) with minor modifications. PTCs or early passages of HEK-293 cells were maintained in high concentration of glucose (25 mM) for 24 h followed by glucose normalization (5.5 mM) for another 24 h (*via* media changing). In pre aPC-treatment condition (HG(aPC)/NG) aPC (20 nM) was added to the high glucose medium, whereas in post aPC-treatment (HG/NG(aPC) aPC (20 nM) was added upon returning cells to normal glucose (5.5 mM glucose).

In a subset of experiments, BUMPT cells were maintained in high concentration of glucose (25 mM) for 48 h with or without SGLT2i (Dapagliflozin®, 2 µM). Other cells were maintained in high concentration of glucose (25 mM) for 24 h followed by glucose normalization (5.5 mM) for another 24 h (*via* media changing).

### *In vitro* Knockdown

Knockdown of protease activated receptors (PARs) or endothelial protein C receptor (EPCR) in PTCs was achieved by lentiviral transduction of short hairpin RNA (shRNA) constructs (pGIPZ-F2r, pGIPZ-F2rl1, pGIPZ-F2rl3, and pGIPZ-Procr, obtained from Dharmacon, Lafayette, CO, USA; targeting mouse PAR1: 5’-ATA TAA GAA GTG ACA TCC A-3’; mouse PAR2: 5’-TTG AGC TGA AGA GTA GGA GC-3’, mouse PAR4: 5’-AAA CAG AGT CCA GTA GTG AGG-3’; and mouse EPCR: 5’-AAA TTC CTG CAG TTC ATA CCG-3’, respectively). A scrambled non-silencing RNA (5’-TAC GAG TAT GAT GTT GGT GGG-3’) was transduced into PTCs as control. Lentiviral particles were generated and concentrated from HEK-293T cells as previously described(*72*) with minor modifications. HEK-293T cells were transduced with pGIPZ-F2r, pGIPZ-F2rl1, pGIPZ-F2rl3 or pGIPZ-Procr together with the packaging plasmid psPAX2 and VSV-G expressing plasmid (pMD2.g; Addgene, USA). The lentiviral particles were harvested from the supernatant 36 and 48 h post-transduction. The lentiviral supernatant was concentrated as previously described(*76*) with minor modifications. The virus-containing medium was mixed with 50% PEG6000 solution, 4 M sodium chloride and 1 x PBS at 26%, 11%, 12% by volume and incubated on a shaker at 4°C for 3-4 h. Following incubation, the solution was aliquoted into 50 ml falcons and centrifuged at the 4700 g at 4°C for 30 min. After centrifugation, the supernatant was carefully removed, and the tube was placed on tissue paper for 1 minute. Cell culture media was added to the semi-dried tube for re-suspension, and then the tube was placed in the 4°C fridge with a cover for overnight recovery. The enriched lentivirus was harvested and aliquoted before being snap frozen. Knockdown efficiency was confirmed by immunoblotting.

### Preparation of activated protein C

Activated protein C (aPC) was generated as previously described (*35, 77*). Prothrombin complex (Prothromplex NF600), containing all vitamin K dependent coagulation factors, was reconstituted with sterile water and supplemented with CaCl2 at a final concentration of 20 mM. The column for purification of protein C was equilibrated at RT with 1 liter of washing buffer (0.1 M NaCl, 20 mM Tris, pH 7.5, 5 mM benzamidine HCl, 2 mM Ca2+, 0.02% sodium azide). The reconstituted prothombin complex was gravity eluted on a column filled with Affigel-10 resin covalently linked to a calcium-dependent monoclonal antibody to PC (HPC4). The column was washed first with two column volumes of washing buffer and then two column volumes with a wash buffer rich in salt (0.5 M NaCl, 20 mM Tris, pH 7.5, 5 mM benzamidine HCl, 2 mM Ca2+, 0.02% sodium azide). Then the benzamidine was washed off the column with a buffer of 0.1 M NaCl, 20 mM Tris pH 7.5, 2 mM Ca2+, 0.02% sodium azide. To elute PC the column was gravity eluted with elution buffer (0.1 M NaCl, 20 mMTris, pH 7.5, 5 mM EDTA, 0.02% sodium azide, pH 7.5) and 3 ml fractions were collected. The peak fractions were identified by measuring absorbance at 280 nm. The peak fractions were pooled. The recovered PC was activated with human plasma thrombin (5% w/w, 3 h at 37°C). To isolate activated protein C (aPC) ion exchange chromatography with FPLC (ÄKTAFPLC®, GE Healthcare Life Sciences) was used. First, thrombin was removed with a cation exchange column MonoS (GE Healthcare Life Sciences). Then a MonoQ anion exchange column (GE Healthcare Life Sciences) was equilibrated with 10% of a 20 mM Tris, pH 7.5, 1 M NaCl buffer. After applying the solution that contains aPC a 10-100% gradient of a 20 mM Tris, pH 7.5, 1 M NaCl buffer was run through the column to elute aPC at a flow of 1-2 ml/min under continuous monitoring of OD and conductivity. APC eluted at ∼36 mS cm-1 by conductivity or at 40% of the buffer. Fractions of 0.5 ml were collected during the peak and pooled. Proteolytic activity of purified aPC was ascertained with the chromogenic substrate SPECTROZYME® PCa and purity was ascertained by a Coomassie stained gel and immunoblotting.

### Human renal biopsies

Human renal biopsy samples were provided by the tissue bank of the National Center for Tumor Diseases (NCT, Heidelberg, Germany) in accordance with the regulations of the tissue bank and the approval of the ethics committee of the University of Heidelberg. For patients’ characteristics see **supplementary table 1**.

### Urine collection and processing

Mouse urine was collected in metabolic cages at indicated time points. Mice were individually placed in metabolic cages to collect 12 h urine samples. Mouse urine was used for determination of albumin and creatinine.

Human urine samples obtained from the local outpatient clinic (patient characteristics in **supplementary table 2**) were obtained based on informed consent (Ethic vote no: 082-10-19042010, University of Leipzig). All participants of the LIFE-ADULT cohort (Ethic vote no: 263-2009-14122009 and 201/17-ek, University of Leipzig, participants characteristics in **supplementary table 3**) (*27*) were fasted on average for 12.8±1.9 h. Urine samples from the FMD study (patient characteristics in **supplementary table 4**) were obtained after ethical approval (Ethic vote no: 204/2004 and 682/2016, Ruprecht-Karls-University of Heidelberg) and informed consent. Urine samples from the LIFE-ADULT cohort were centrifuged at 2750 g for 10 min within 2h and stored at –80°C by the LIFE-Biobank team, part of the Leipzig Medical Biobank. Other urine samples were centrifuged at 5000 g for 5 min to remove debris and dead cells. Human urine supernatants were used for determination of p21 by immunoblotting or ELISA (see next section). For p21 immunoblotting, aliquots of the human urine supernatant were centrifuged again at 20000 g for 60 min and the supernatant was mixed with laemmli sample buffer containing protease inhibitors.

### p21 ELISA for human urine samples

For determination of urinary p21 by ELISA we used anti-human p21 matched antibody pair Kit (abcam, USA) according to manufacturer’s instructions. Maxisorb plates were coated with p21 capture antibody overnight at 4°C followed by washing and blocking at 37°C for 1 h. Human urine supernatants were diluted (1:2), added to the microplates and incubated for 2 h at room temperature on a plate shaker set to 400 rpm. This was followed by washing (3x, using wash buffer) and incubation with detector antibody for 1 h at room temperature on a plate shaker at 400 rpm. Following washing (3x, using wash buffer), HRP-Streptavidin solution was added and the plate was incubated for 1 h at room temperature on a plate shaker at 400 rpm. Plates were washed (3x using wash buffer) and tetramethylbenzidine (TMB) substrate was added for color development. Reaction was stopped by adding stop solution (as provided in the kit) and absorbance was measured at 450 nm after 10 min.

### Albuminuria in mouse urine samples

Mouse urine albumin and creatinine were measured as previously described(*13, 72*, 78). Urine albumin was determined using a mouse albumin ELISA kit (Mouse albumin ELISA quantification kit, Bethyl Laboratories) according to the manufacturer’s instructions. Urine creatinine was determined using a commercially available assay (Roche Diagnostics, Cobas c501 module) in the Institute of Clinical Chemistry and Pathobiochemistry, Medical Faculty, Otto-von-Guericke University, Magdeburg, Germany and institute of laboratory medicine, clinical chemistry and molecular diagnostics, Leipzig, Germany(*35*).

### Methylation specific PCR (MSP)

Methylation-specific PCR was performed as previously described (*9, 11*). Genomic DNA was extracted from HEK-293 cells or kidney tissues using the QIAamp DNA Mini kit (QIAGEN) and treated with sodium bisulfite using the EZ DNA Methylation kit (Zymo Research) according to the manufacturer’s instructions. PCR amplification was performed with the EpiTect MSP kit (QIAGEN) using primer pairs designed to specifically detect either methylated or unmethylated CpG sites in the human or mouse p21 promoter: human methylated forward, 5′-TTT GTT GTA TGA TTT GAG TTA G-3′; human methylated reverse, 5′-TAA TCC CTC ACT AAA TCA CCT C-3′; human unmethylated forward, 5′-ATC ATT CTG GCC TCA AGA TGC-3′; and human unmethylated reverse, 5′-CGG CTC CAC AAG GAA CTG AC-3′; mouse methylated forward 5′-GTT AGC GAG TTT TCG GGA TC-3′; mouse methylated reverse 5′-CTC GAC TAC TAC AAT TAA CGT CGA A-3′; mouse unmethylated forward 5′-GGT TAG TGA GTT TTT GGG ATT G-3′; mouse unmethylated reverse 5′-TCT CAA CTA CTA CAA TTA ACA TCA AA-3′. For ratio analyses, amplicons against methylated and unmethylated CpGs were visualized on a 2% (w/v) agarose gel. Universal methylated human and mouse DNA (Millipore) was used as a positive control.

### DNMT activity assay

DNMTs activity was measured by EpiQuik DNA Methyltransferase Activity/Inhibition assay kit (Epigentek) according to the manufacturer’s instructions. Confluent HEK-293 cells were treated with a cytoplasmic extraction buffer (10 mM HEPES, 1.5 mM MgCl_2_, 10 mM KCl and 0.5 mM DTT), briefly vortexed and then centrifuged (14000 g, 4°C, 30 sec). Supernatants containing the cytoplasmic protein fractions were discarded. The pellet was re-suspended in 100 μl of a nuclear extraction buffer (20 mM HEPES, 420 mM NaCl, 1.5 mM MgCl_2_, 0.2 mM EDTA, 0.5 mM DTT and 25% glycerol), incubated for 20 min on ice, followed by centrifugation (13.000 g, 4°C, 5 min). Supernatants containing DNMTs within the nuclear extracts were collected and added to wells coated with cytosine-rich DNA-substrate and pre-incubated with the DNMT assay buffer and the methyl group donor Adomet. Wells were covered and incubated for 1 h at 37°C. Wells were then washed three times and the capture antibody against 5-methylcytosine was added. After 1 h incubation on an orbital shaker at room temperature wells were washed 4 times, the secondary detection antibody was added, and samples were incubated for 30 min at room temperature. Wells were washed five times and developer solution was added for 5 min before adding the stop solution. Absorbance was immediately measured at 450 nm.

### Immunoblotting

Immunoblotting was conducted as previously described (*13, 72*). Cell lysates were prepared using RIPA buffer containing 50 mM Tris (pH7.4), 1% NP-40, 0.25% sodium-deoxycholate, 150 mM NaCl, 1 mM EDTA, 1 mM Na_3_VO_4_, 1 mM NaF supplemented with protease inhibitor cocktail. Lysates were centrifuged (13000 g for 10 min at 4°C) and insoluble debris was discarded. Protein concentration in supernatants was quantified using BCA reagent. Equal amounts of protein were electrophoretically separated on 7.5%, 10% or 12.5% SDS polyacrylamide gel, transferred to PVDF membranes and probed with desired primary antibodies at appropriate dilution (KIM-1: 1:2000; SC-p21: 1:350; CST-p21: 1:1000; DNMT1: 1:1000; GAPDH: 1:25000; β-actin: 1:1000). After overnight incubation with primary antibodies at 4°C membranes were washed with TBST and incubated with anti-rabbit IgG (1:2000) horseradish peroxidase-conjugated antibody for 1 hr at room temperature. Blots were developed with the enhanced chemiluminescence system. To compare and quantify levels of proteins, the density of each band was measured using ImageJ software. Equal loading for total cell or tissue lysates was determined by GAPDH or β-actin immunoblots using the same blot whenever possible.

### Reverse transcriptase PCR (RT-PCR)

RT-PCR was conducted essentially as previously described with modifications (*9, 72*). Kidney tissues stored in RNA*later*™ (Invitrogen) were thawed on ice and transferred directly into Trizol (Life Technologies) for isolation of total RNA following the manufacturer’s protocol. For *in viro* cells, Trizol was added to the cells after removing the medium and washing with PBS. Quality of total RNA was ensured on agarose gel and by analysis of the A260/280 ratio. The reverse transcription reaction was conducted using 1 μg of total RNA after treatment with DNase I (5 U/5 μg RNA, 30 min, 37°C) followed by reverse transcription using RevertAid First Strand cDNA Synthesis Kit (Thermo Fisher Scientific). Semi-quantitative polymerase chain reactions were performed and gene expression was normalized to β-actin. PCR products were separated on a 1.5 – 1.8% agarose gel and visualized by ethidium bromide (Et-Br) staining. Reactions lacking reverse transcription served as negative controls. For primers sequence see **supplementary table 5**.

### Quantitative real time PCR (qRT-PCR)

Total RNA was extracted from kidney tissues, RNA quality was ensured, and cDNA was generated as outlined in the previous section. qRT-PCR was carried out on CFX Connect Real-time system (Bio-Rad Laboratories, Hercules, CA) using SYBR Green (Takyon™, Eurogentec). For quantitative analysis results were normalized to 18srRNA. Gene expression was analyzed using the 2^−(ΔΔCt)^ method(*79*). Reactions lacking cDNA served as negative controls. For primers sequence see **supplementary table 5**.

### RNA expression profiling

RNA was isolated from murine renal cortex using RNAeasy mini kit (QIAGEN). RNA was isolated from non-diabetic, age matched control mice (C, C57Bl/6), from mice with 22 weeks of persistent hyperglycemic following STZ injection as outlined above (DM), or from mice with 16 weeks of persistent hyperglycemia followed by 6 additional weeks with lowered blood glucose levels using SGLT2i (DM+SGLT2i). In another set of experiments, we isolated RNA from aged matched (22 weeks age) normoglycemic control mice (db/m), hyperglycemic diabetic mice (db/db), and db/db mice with initially elevated, but then markedly reduced blood glucose levels for 6 weeks prior to analyses (db/db+SGLT2i). Quality and integrity of total RNA was verified using an Agilent Technologies 2100 Bioanalyzer (Agilent Technologies; Waldbronn, Germany).

For expression profiling by RNA sequencing (RNA-seq) a RNA sequencing library was generated from 500 ng total RNA. The Dynabeads® mRNA DIRECT™ Micro Purification Kit (Thermo Fisher) was used for purification of mRNA. The RNA sequencing library was generatd using the ScriptSeq v2 RNA-Seq Library Preparation Kit (Epicentre) according to manufacturer’s protocols. The libraries were sequenced on Illumina HiSeq2500 using TruSeq SBS Kit v3-HS (50 cycles, single ended run) with an average of 3 ×10^7^ reads per RNA sample. For each FASTQ file a quality report was generated by FASTQC tool. Before alignment to reference genome each sequence in the raw FASTQ files were trimmed on base call quality and sequencing adapter contamination using Trim Galore! wrapper tool. Reads shorter than 20 bp were removed from FASTQ files. Trimmed reads were aligned to the reference genome using open source short read aligner STAR (https://code.google.com/p/rna-star/) with settings according to log file. Feature counts were determined using R package “Rsubread”. Only genes showing counts greater than 5 at least two times across all samples were considered for further analysis (data cleansing). Gene annotation was done by R package “bioMaRt”. Before starting the statistical analysis steps, expression data was log2 transformed and normalized according to the 50^th^ percentile (quartile normalization using edgeR). Differential gene expression was calculated by R package “edgeR”. The statistical analysis was performed on the filtered and annotated data with Limma package (moderated t statistics). To correct for multiple testing error, the Benjamini-Hochberg correction (BH) was used.

### Functional annotation and Pathway analysis

The threshold to identify differentially expressed genes (DEGs) was set to a logFc value of ±0.58 resulting in 1.5-fold expression change. Statistically significant DEGs (p<0.01; FDR<0.1) were categorized into genes with persistently changed expression despite glucose lowering and genes with normalized expression upon glucose lowering. Heatmapper (heatmapper.ca) was used to generate heatmaps of gene expression data. Row dendrograms were generated using hierarchical row clustering calculated by squared Euclidean distances (*80*). pantherDB (http://www.pantherdb.org/) was used for pathway analysis. The percentage of gene hits relative to total number of pathway hits was used to evaluate prominent pathways involved.

### Histology, immunohistochemistry and histological analyses

Tubular morphometrics were determined using a histological score based on periodic acid Schiff’s (PAS) stained images, following an established protocol (*81*). At least 50 glomerular and tubular cross sections were analyzed for each mouse. For determination of the tubular and glomerular parameters adjacent sections were compared to identify the maximal glomerular diameter and to ensure that a transversal tubular cross-sections were analyzed and to avoid artefacts from tangentially cut glomeruli or tubuli. To calculate the epithelial surface area of tubular cross sections, the difference of the area of external tubuli profile and the area of the tubular lumen was determined (*81*). Tubular cell hyperplasia was evaluated on the same tubular cross sections by counting the number of nuclei(*81*). Tubular cell hypertrophy was calculated as the ratio between epithelial surface and the number of nuclei for tubular section (*81*).

Renal tubulointerstitial fibrosis was evaluated using Masson’s trichrome staining (MTS), as previously described (*72, 82*). Ten visual fields per slide were randomly selected at 200 x magnification. Interstitial fibrosis was quantified using NIH-ImageJ software, generating a binary image allowing automatic calculation of the stained area as percentage of the image area (*72, 82*). The glomerular fractional mesangial area (FMA) was calculated following the current DCC (Diabetes Complications Consortium) protocol and as described before (*72*). In brief, 5 μm thick sections were stained with PAS reagent. In every investigated glomerulus, the tuft glomerular area was determined using NIH-ImageJ software. FMA was calculated as the fractional area positive for PAS in the glomerular tuft area, reported as the percentage.

p21 and KIM-1 expression were determined by immunohistochemistry (IHC) following established protocols (*9, 83*). The number of p21-positive cells (defined as those stained brown using a DAB substrate, 3,3’-diaminobenzidin; hematoxylin counterstain) was determined in at least 10 randomly chosen fields per section at a magnification of 40 x in the renal cortical area (*42*).

Images for all histological analyses were obtained using an Olympus BX43 microscope, Olympus XC30 Camera, and Olympus cellSens Dimension 1.5 Image software. All images of a specific analytical method (e.g. FMA, IHC-stains) were taken with the same settings. The evaluations were performed by a blinded investigator.

### Senescence associated beta galactosidase (SA-β-gal.) staining

Renal cell senescence was evaluated using Senescence Cells Histochemical Staining Kit (CS0030-Sigma) according to the manufacturer’s protocol. 5 μm thick cryro-sections of renal tissue or early passage HEK-293 cells were washed in 1 x PBS and fixed for 1 min (2% formaldehyde, 0.2% glutaraldehyde, 7.04 mM Na_2_HPO_4_, 1.47 mM KH_2_PO_4_, 0.137 M NaCl, 2.68 mM KCl) and then rinsed in ice-cold 1 x PBS for 5 min. The cryo-sections were incubated with freshly prepared X-gal staining solution (10 ml containing 0.25 ml of 40 mg/ml X-gal Solution, 125 µl of 400 mM potassium ferricyanide, 125 µl of 400 mM potassium ferrocyanide) at 37°C overnight. Next, excess staining solution was washed away using ice-cold 1 x PBS, and the sections were overlaid with 70% glycerol followed by staining assessment. For frozen sections, aqueous Eosin was used for counterstaining before mounting the slides with 70% glycerol. The development of blue dots/areas was counted as positive staining (*42, 84*). Senescence was evaluated as the ratio (in percent) of the blue-colored area in each sample relative to the total area of at least 10 renal cortical visual fields per section (*42*).

### Statistical Analysis

The data are summarized as mean ± SEM (standard error of the mean). Statistical analyses were performed with parametric Student’s t-test and ANOVA or non-parametric Mann-Whitney-Test and Kruskal-Wallis test, as appropriate, and post-hoc comparison with the method of Tukey, Bonferroni or Sidak’s multiple comparison. The Kolmogorov-Smirnov (KS) test or D’Agostino-Pearson-Normality-test was used to determine whether the data are consistent with a Gaussian distribution. StatistiXL (www.statistixl.com) and Prism 5 (www.graphpad.com) softwares were used for statistical analyses. Statistical significance was accepted at values of p<0.05.

## Acknowledgement

We thank Kathrin Deneser, Julia Judin, Juliane Friedrich, René Rudat, and Rumiya Makarova for excellent technical support. This work was supported by grants of the ‘Deutsche Forschungsgemeinschaft’ (DFG, German Research Foundation: IS-67/8-1, IS-67/11-1, SFB854/B26, RTG2408/P7&P9 to B.I., SFB854/A01, ME-1365/7-2, ME1365/9-2 to P.R.M., RTG2408/P5, SH 849/1-2 to K.S., KO 5736/1-1 to S.K., and Projektnummer 236360313 – SFB 1118 to BI and PPN), of the ‘Stiftung Pathobiochemie und Molekulare Diagnostik’ (SPMD to K.S.), by DAAD scholarships to M.M.A. and A.E., and by LIFE – Leipzig Research Centre for Civilization Diseases, an organizational unit affiliated to the Medical Faculty of the University of Leipzig. LIFE is funded by means of the European Union, by the European Regional Development Fund (ERDF), by funds of the Free State of Saxony within the framework of the excellence initiative (project numbers 713-241202, 713-241202, 14505/2470, 14575/2470), and by funds of the Medical Faculty of the University of Leipzig.

## Author Contribution

M.M.A, K.S., A.E., and S.Ko. designed, performed, and interpreted in vivo, in vitro, and ex vivo experiments; A.S., S.K., S.Z., P.M., and P.P.N. provided human samples and assisted in preparing the manuscript, F.B., I.G., S.N., R.R., S.Kr., D.G. assisted in performing in vivo studies, ex vivo analysis and histological analyses; R.G. conducted expression analyses and assisted in data interpretation, J.G. and C.D. provided reagents and interpreted data, S.Ko. conducted functional annotation, data analyses, in silico studies, and assisted in preparing the manuscript, B.I. conceptually designed and interpreted the experimental work and prepared the manuscript.

## Materials and Correspondence

For general comments or material requests please contact B.I.

## Data availability

The data that support the findings of this study are available from the corresponding author upon reasonable request. Sequencing data that support the findings of this study have been deposited at an FTP server (genome.gbf.de).

