## Supplemental tables for "Reversal of the renal hyperglycemic memory by targeting sustained tubular p21 expression"

*Running headline: Reversing hyperglycemic memory in DKD*

[1] Institute of Laboratory Medicine, Clinical Chemistry and Molecular Diagnostics, Universitätsklinikum Leipzig, Leipzig University, Leipzig, Germany.
[2] Department of Medical Laboratories, Faculty of Health Sciences, American University of Madaba (AUM), Amman, Jordan.
[3] Department of Biotechnology, University of Sargodha, Pakistan.
[4] Internal Medicine I and Clinical Chemistry, German Diabetes Center (DZD), University of Heidelberg, Heidelberg, Germany.
[5] Department of Medicine, Vanderbilt University Medical Center, Nashville, Tennessee, United States.
[6] Leipzig Medical Biobank, Leipzig University, Leipzig, Germany.
[7] Institute for Medical Informatics, Statistics and Epidemiology, Leipzig University, Leipzig, Germany.
[8] Helmholtz Centre for Infection Research, Braunschweig, Germany.
[9] Clinic of Nephrology and Hypertension, Diabetes and Endocrinology, Otto-von-Guericke University, Magdeburg, Germany.
[10] Department of Molecular Medicine, The Scripps Research Institute, La Jolla, CA, USA.
[11] Department of Chemistry, Marquette University, Milwaukee, WI, USA.

*These authors contributed equally to the work.

| Characteristic | C (n=6) | DM (-DKD) (n=6) | DM (+DKD) (n=5) |
|---|---|---|---|
| **Age** (years) | 71.5 ± 3.1 | 75.0 ± 3.2 | 78.2 ± 2.5 |
| **Sex** (female/male) | (3/3) | (3/3) | (2/3) |
| **History (N) of** | | | |
| Hypertension | 2 | 4 | 4 |
| Brain tumor | 1 | - | - |
| COPD | 1 | - | - |
| Steatosis hepatitis | 1 | - | - |
| peripheral arterial occlusive disease | 1 | - | - |
| Hypothyroidism | 1 | - | - |
| Type 2 diabetes | - | 6 | 5 |
| **Diabetic complications (N)** | | | |
| Diabetic neuropathy | - | 2 | - |
| Diabetic kidney disease (DKD) | - | - | 5 |
| **Serum creatinine** (mg/dl) | 1.1 ± 0.17 | 1.1 ± 0.06 | 1.9 ± 0.36 |
| **Serum urea** (mg/dl) | 6.6 ± 0.66 | 6.8 ± 0.69 | 10.8 ± 1.78 |

**Supplementary Table 1 (corresponding to Fig. 2a-c). Anthropometric, clinical and metabolic characteristics of renal biopsy control and diabetic patients with and without diabetic kidney disease.** Data represents mean ± SEM. Abbreviations: C: Control, DKD: Diabetic Kidney Disease, DM: Diabetes Mellitus, COPD: chronic obstructive pulmonary disease.

| Characteristic | Control Baseline (n=22) | DKD Baseline (n=26) |
|---|---|---|
| **Age** (years) | 64.2 ± 2.1 | 65 ± 2.6 **n.s.** |
| **Sex** (female/male) | (8/14) | (5/21) |
| **Diabetes complication** | | |
| Nephropathy (n) | - | 26 |
| Retinopathy (n) | - | 3 |
| Neuropathy (n) | - | 10 |
| **History of** | | |
| Hypertension (n) | 7 | 18 |
| Coronary heart disease | 0 | 12 |
| Myocardial infarction (n) | 0 | 4 |
| OSAS (n) | 3 | 3 |
| Arthrosis (n) | 5 | 4 |
| Thyroid nodules (n) | 0 | 1 |
| Thyreoidectomy (n) | 2 | 0 |
| **Diabetes therapy** | | |
| Metformin (n) | - | 5 |
| DPP-4 (n) | - | 4 |
| SGLT-2 (n) | - | 1 |
| Glinide (n) | - | 0 |
| Long-acting insulin | - | 18 |
| **Other medication** | | |
| RAAS-inhibitors (n) | 5 | 12 |
| Beta-blockers (n) | 2 | 15 |
| Thiazide diuretics (n) | 0 | 2 |
| Loop diuretics (n) | 2 | 13 |
| Calcium-antagonist (n) | 1 | 5 |
| Statin (n) | 2 | 17 |
| Ezetimib (n) | 0 | 4 |
| Acetyl-salicylic acid (n) | 0 | 7 |
| L-Thyroxin | 3 | 3 |
| **Glycemic control** | | |
| HbA1c (%) | 5.7 ± 0.2 | 7.5 ± 0.2** |
| RPG (mmol/l) | 6.1± 0.8 | 9.9 ± 1.1* |
| **Serum creatinine** (mg/dl) | 0.86 ± 0.17 | 1.67 ± 0.71*** |
| **eGFR** (ml/min/1.73m²) | 77,6 ± 3,5 | 43,9 ± 5,6*** |
| **Albuminuria** (mg albumin/g urine-creatinine) | 7,2 ± 0,8 | 537,4 ± 152,9** |

**Supplementary Table 2 (corresponding to Fig. 2e). Clinical and metabolic characteristics of non-diabetic controls and diabetic patients with known DKD, both recruited from the local outpatient clinic.** Data represents mean ± SEM, comparison by t-test, **P*<0.05, ** *P*<0.01, *** *P*<0.001 and n.s: non-significant. Abbreviations: DPP-4: dipeptydil peptidase 4, RAAS: renin-angiotensin-aldosterone system, RBG: random plasma glucose, eGFR: estimated glomerular filtration rate.

| Characteristic | Control | Low risk | Moderate risk | High risk | Very high risk |
|---|---|---|---|---|---|
| **Number of patients** | **n=36** | **n=52** | **n=53** | **n=29** | **n=18** |
| **Age** (years) | (63.2 ± 0.4) | (71.0 ± 0.6)*** | (70.1 ± 0.7)*** | (71.0 ± 0.9)*** | (74.1 ± 1.1)*** |
| **Sex** (female/male) | (15/22) | (7/45) | (15/38) | (11/18) | (5/13) |
| **BMI** (kg/m²) | 28.38±0.76 | 30.1±4.6 | 30.7±4.5* | 30.6±4.3* | 31.2±4.3* |
| **Blood pressure** | | | | | |
| systolic (mmHg) | 135.28 ± 2.91 | 138.8 ± 19.2 | 131.8 ± 18.3 | 134.1 ± 19.8 | 147.6 ± 32.4 |
| diastolic (mmHg) | 76.92 ± 1.73 | 73.9 ± 9.2 | 71.1 ± 8.9** | 70.6 ± 8.6** | 77.7 ± 14.0 |
| **Diabetes therapy (N)** | - | | | | |
| Metformin | - | 28 | 37 | 18 | 5 |
| DPP-4 | - | 8 | 17 | 7 | 6 |
| Glinide | - | 6 | 7 | 7 | 4 |
| Long-acting insulin | - | 8 | 14 | 11 | 12 |
| **Other medication (N)** | | | | | |
| RAAS-inhibitors | 6 | 37 | 41 | 20 | 18 |
| Beta-blockers | 5 | 23 | 27 | 15 | 12 |
| Thiazide diuretics | 0 | 8 | 0 | 0 | 0 |
| Loop diuretics | 0 | 2 | 0 | 0 | 0 |
| Calcium-antagonist | 0 | 9 | 19 | 20 | 6 |
| Statin | 1 | 27 | 23 | 14 | 7 |
| Ezetimib | 0 | 2 | 0 | 0 | 0 |
| Acetyl-salicylic acid | 1 | 17 | 0 | 0 | 0 |
| L-Thyroxin | 3 | 7 | 11 | 3 | 1 |
| **Glycemic control** | | | | | |
| HbA1c (%) | 5.40 ± 0.10 | 6.1 ± 0.7*** | 6.4 ± 0.7*** | 6.2 ± 1.2** | 6.0 ± 0.6** |
| **Serum creatinine** (mg/dl) | (0.99 ± 0.10) | (0.92 ± 0.02) | (0.95 ± 0.02) | (1.05 ± 0.05) | (1.50 ± 0.07)*** |
| **eGFR** (ml/min/1.73m²) | (99.4 ± 0.4) | (79.8 ± 1.3)*** | (68.9 ± 1.4)** | (73.9 ± 3.2)*** | (40.9 ± 3.0)*** |
| **Albuminuria** (mg albumin/g urine-creatinine) | (8.8 ± 1.12) | (11.1 ± 1.0) n.s. | (85.4 ± 10.4)*** | (730 ± 203.8)*** | (802 ± 176.2)*** |

**Supplementary Table 3 (corresponding to Fig. 2f-j). Anthropometric, clinical and metabolic characteristics of non-diabetic controls and diabetic individuals from the LIFE-adult cohort.** The severity of DKD was classified according to the KDIGO criteria as low, moderate, high and very high risk of chronic kidney disease (CKD). Data represents mean ± SEM, comparison by t-test (each diabetic group against controls), *$P$<0.05, ** $P$<0.01, *** $P$<0.001 and n.s: non-significant. Abbreviations: BMI: body mass index, DPP-4: dipeptydil peptidase 4, RAAS: renin-angiotensin-aldosterone system, eGFR: estimated glomerular filtration rate.

| Characteristic | Baseline (n=10) | After 3 cycles FMD (n=10) |
|---|---|---|
| **Age** (years) | 64.8 ± 2.4 | -- |
| **Sex** (female/male) | 4/10 | -- |
| **BMI** (kg/m²) | 30.4 ± 1.6 | 28.5 ± 1.6 |
| **Diabetes duration** (years) | 15.0 ± 2.2 | -- |
| **Diabetes complication (n)** | | |
| Nephropathy | 10 | 8 |
| Retinopathy | 2 | 2 |
| Neuropathy | 4 | 4 |
| **History (n) of** | | |
| Hypertension | 10 | 10 |
| Coronary heart disease | 2 | 2 |
| Myocardial infarction | 1 | 1 |
| OSAS | 1 | 1 |
| Arthrosis | 4 | 4 |
| Thyroid nodules | 2 | 2 |
| Thyreoidectomy | 2 | 2 |
| **Diabetes therapy (n)** | | |
| Metformin | 9 | 9 |
| DPP-4 | 4 | 3 |
| SGLT2 inhibitors | 3 | 1 |
| Glinide | 1 | 0 |
| Long-acting insulin | 1 | 1 |
| **Other medication (n)** | | |
| RAAS-inhibitors | 9 | 9 |
| Beta-blockers | 5 | 5 |
| Thiazide diuretics | 4 | 4 |
| Loop diuretics | 3 | 3 |
| Calcium-antagonist | 3 | 4 |
| Statin | 3 | 3 |
| Ezetimib | 2 | 2 |
| Acetyl-salicylic acid | 2 | 3 |
| **Glycemic control** | | |
| HbA1c (%) | 7.4 ± 0.3 | 6.9 ± 0.3** |
| FPG (mg/dl) | 8.6 ± 0.7 | 7.8 ± 1.1 |
| **Blood pressure** (systolic/diastolic mmHg) | 138.6/83.3 ± 3.8/3.0 | 137.4/80.7 ± 3.3/2.7 |
| **Serum creatinine** (mg/dl) | 0.9 ± 0.1 | 0.9 ± 0.1 |
| **eGFR** (ml/min/1.73m²) | 93.1 ± 7.1 | 96.9 ± 7.7 |
| **eGFR from cystatin C** (ml/min/1.73m²) | 88.2 ± 5.9 | 86.8 ± 5.4 |
| **Total cholesterol** (mg/dl) | 184.2 ± 12.9 | 167.0 ± 13.4 |
| **LDL** (mg/dl) | 101.5 ± 12.7 | 97.2 ± 13.4 |
| **HDL** (mg/dl) | 45.4 ± 3.2 | 44.5 ± 3.7 |
| **Triglycerides** (mg/dl) | 245.5 ± 77.3 | 125.7 ± 20.7 |
| **Albuminuria** (mg albumin/g urine-creatinine) | 62.8 ± 16.2 | 47.9 ± 8.7 |

**Supplementary Table 4 (corresponding to Fig. 2k-m). Anthropometric, clinical and metabolic characteristics of diabetic patients before and after 3 cycles of FMD (fasting mimicking diet).** Data represents mean ± SEM, comparison by t-test, ***P*<0.01. Abbreviations: BMI: body mass index, OSAS: obstructive sleep apnea syndrome, DPP-4: dipeptydil peptidase 4, SGLT2: sodium-glucose cotransporter 2, RAAS: renin-angiotensin-aldosterone system, FPG: fasting plasma glucose, eGFR: estimated glomerular filtration rate, LDL: low-density Lipoprotein, HDL: high-density lipoprotein. The study is registered in the German Clinical Trials Register (DRKS00014287).

| Target | Sequence |
| --- | --- |
| Mt-hp21 | 5’-TTT GTT GTA TGA TTT GAG TTA G-3‘ |
| | 5’-TAA TCC CTC ACT AAA TCA CCT C-3‘ |
| Um-hp21 | 5’-ATC ATT CTG GCC TCA AGA TGC-3‘ |
| | 5’-CGG CTC CAC AAG GAA CTG AC-3‘ |
| hDNMT1 | 5’-GGC GGC TCA AAG ATT TGG AAA GAG-3‘ |
| | 5’-CAC CGT TCT CCA AGG ACA AAT C-3‘ |
| hDNMT3a | 5’-CTT TGA TGG GAT TGC TAC AGG-3‘ |
| | 5’-ACA CCT CGG AGG CAA TGT AG-3‘ |
| hDNMT3b | 5’-CAG GGA AAA CTG CAA AGC TC-3‘ |
| | 5’-ATT TGT TAC GTC GTG GCT CC-3‘ |
| hp21 | 5’-CCG AAG TCA GTT CCT TGT GG-3‘ |
| | 5’-CAT GGG TTC TGA CGG ACA T-3‘ |
| hβ-actin | 5’-GCC TCG CCT TTG CCG AT-3‘ |
| | 5’-CCA CGA TGG AGG GGA AGA C-3‘ |
| Mt-mp21 | 5’-GTT AGC GAG TTT TCG GGA TC-3‘ |
| | 5’-CTC GAC TAC TAC AAT TAA CGT CGA A-3‘ |
| Um-mp21 | 5’-GGT TAG TGA GTT TTT GGG ATT G-3‘ |
| | 5’-TCT CAA CTA CTA CAA TTA ACA TCA AA-3‘ |
| mp21 | 5’-TCC ACA GCG ATA TCC AGA CA-3‘ |
| | 5’-GGA CAT CAC CAG GAT TGG AC-3‘ |
| mDNMT1 | 5’-CAG AGA CTC CCG AGG ACA GA-3‘ |
| | 5’-TTT ACG TGT CGT TTT TCG TCT C-3‘ |
| mDNMT3a | 5’-CGT GAG TCC GGT GTG TCA-3‘ |
| | 5’-CTC CAA CCA CAC ACA CAA GG-3‘ |
| mDNMT3b | 5’-TTC AGT GAC CAG TCC TCA GAC ACG AA-3‘ |
| | 5’-TCA GAA GGC TGG AGA CCT CCC TCT T-3‘ |
| mKIM-1 | 5’-TCC ACA CAT GTA CCA ACA TCA A-3‘ |
| | 5’-GTC ACA GTG CCA TTC CAG TC-3‘ |
| mPAR1 | 5’-CCA GCC AGA ATC AGA GAG GA-3‘ |
| | 5’-CGG AGA TGA AGG GAG GAG-3‘ |
| mPAR2 | 5’-CCA GGA AGA AGG CAA ACA TC-3‘ |
| | 5’-TGT CCC CCA CCA ATA CCT C-3‘ |
| mPAR3 | 5’-CAT CCT GCT GTT TGT GGT TG-3‘ |
| | 5’-TAC CCA GTT GTT GCC ATT GA-3‘ |
| mPAR4 | 5’-GCA GAC CTT CCG ATT AGC TG-3‘ |
| | 5’-CAC TGC CGA GAA CAG TAC CA-3‘ |
| mEPCR | 5’-CTA CAA CCG GAC TCG GTA TGA A-3‘ |
| | 5’-CCA GGA CCA GTG ATG TGT AAG A-3‘ |
| mβ-actin | 5’-CTA GAC TTC GAG CAG GAG ATG G-3‘ |
| | 5’-GCT AGG AGC CAG AGC AGT AAT C-3‘ |
| 18srRNA | 5’-GGC CCT GTA ATT GGA ATG AGT C-3‘ |
| | 5’-CCA AGA TCC AAC TAC GAG CTT-3‘ |

**Supplementary Table 5:** Primers sequences as used in the current study. Abbreviations: h: human; m: mouse; Mt: methylated; Um: unmethylated
