## Supplemental figures for "Reversal of the renal hyperglycemic memory by targeting sustained tubular p21 expression"

#### Supp. Fig. 1 (related to Fig. 1)

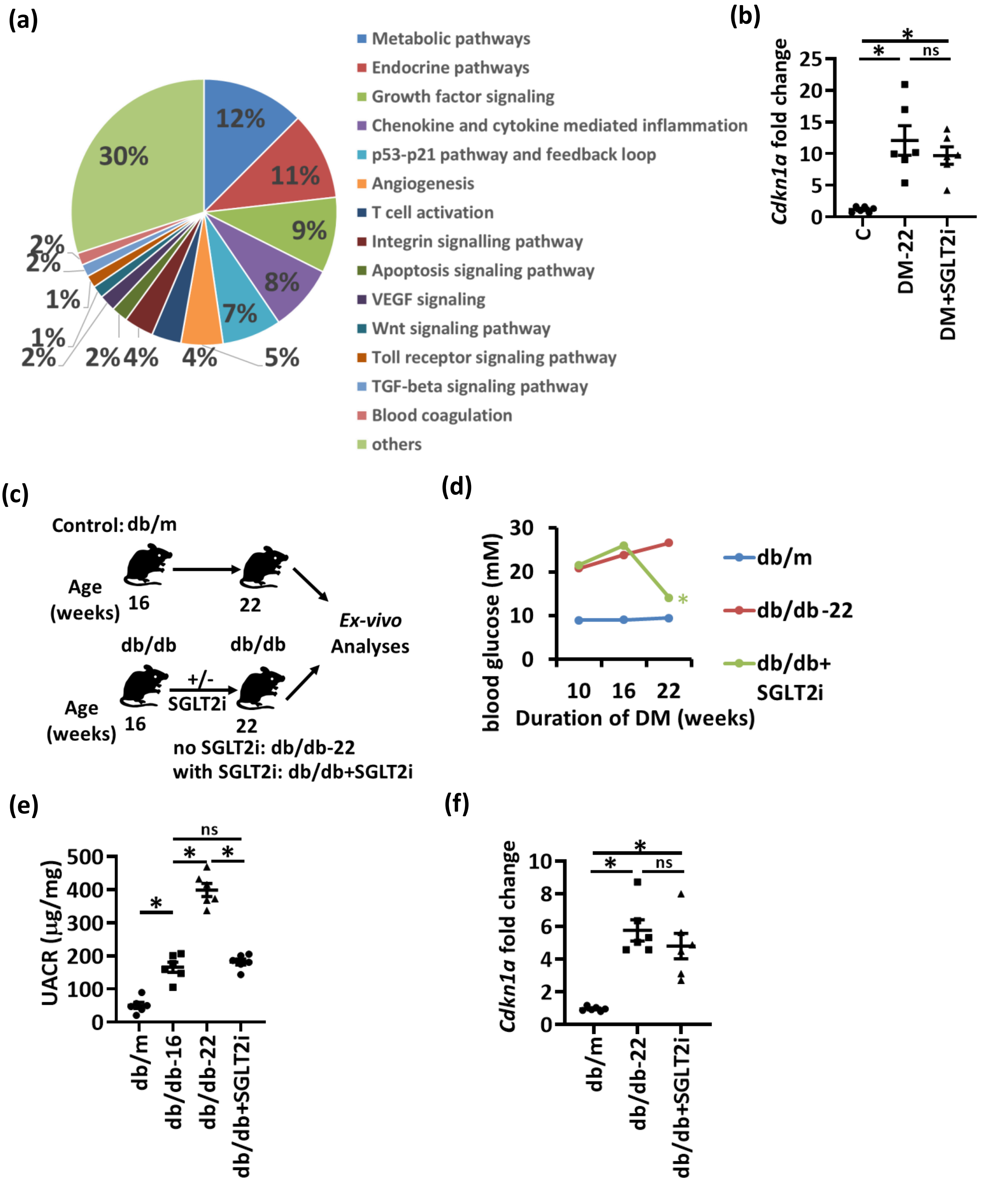

**Supp. Fig. 1: Sustained p21 expression despite reduced blood glucose levels in experimental type 1 and type 2 DM**

**a)** Pie chart reflecting renal pathways identified based on genes persistently changed despite lowering blood glucose in type 1 diabetic mice.
**b)** Dot plot summarizing renal p21 mRNA (*Cdkn1a*, qRT-PCR). Experimental groups: non-diabetic controls (C) and type 1 diabetic mice. 16 weeks after induction of persistent hyperglycemia, diabetic mice were treated with PBS (DM-22) or sodium/glucose cotransporter 2-inhibitor (Dapagliflozin®, SGLT2i; DM+SGLT2i) for further 6 weeks. Dot plot reflecting mean ± SEM of at least 6 mice per group; ANOVA, *$P$<0.05. ; ns: non-significant.
**c)** Experimental scheme of type 2 diabetes mellitus model (T2DM): 16 weeks old db/db mice were treated with PBS (db/db-22) or with the sodium/glucose cotransporter 2-inhibitor (Dapagliflozin®, SGLT2i, db/db+SGLT2i) for further 6 weeks. Non-diabetic mice (db/m) were used as controls.
**d)** Average blood glucose levels in experimental groups (T2DM-model, as described in c) after 10 or 16 weeks of hyperglycemia and at 22 weeks. Line graphs reflecting mean ± SEM of at least 6 mice per group; ANOVA, *P<0.05.
**e)** Dot plot summarizing albuminuria (urinary albumin-creatinine ratio, µg albumin/mg creatinine; UACR) in experimental groups (as described in c). Dot plot reflecting mean ± SEM of at least 6 mice per group; ANOVA, *P<0.05; ns: non-significant.
**f)** Dot plot summarizing renal p21 mRNA (*Cdkn1a*, qRT-PCR) in experimental groups (as described in c). Dot plot reflecting mean ± SEM of at least 6 mice per group; ANOVA, *$P$<0.05. ; ns: non-significant.

**Supp. Fig. 2 (related to Fig. 1)**

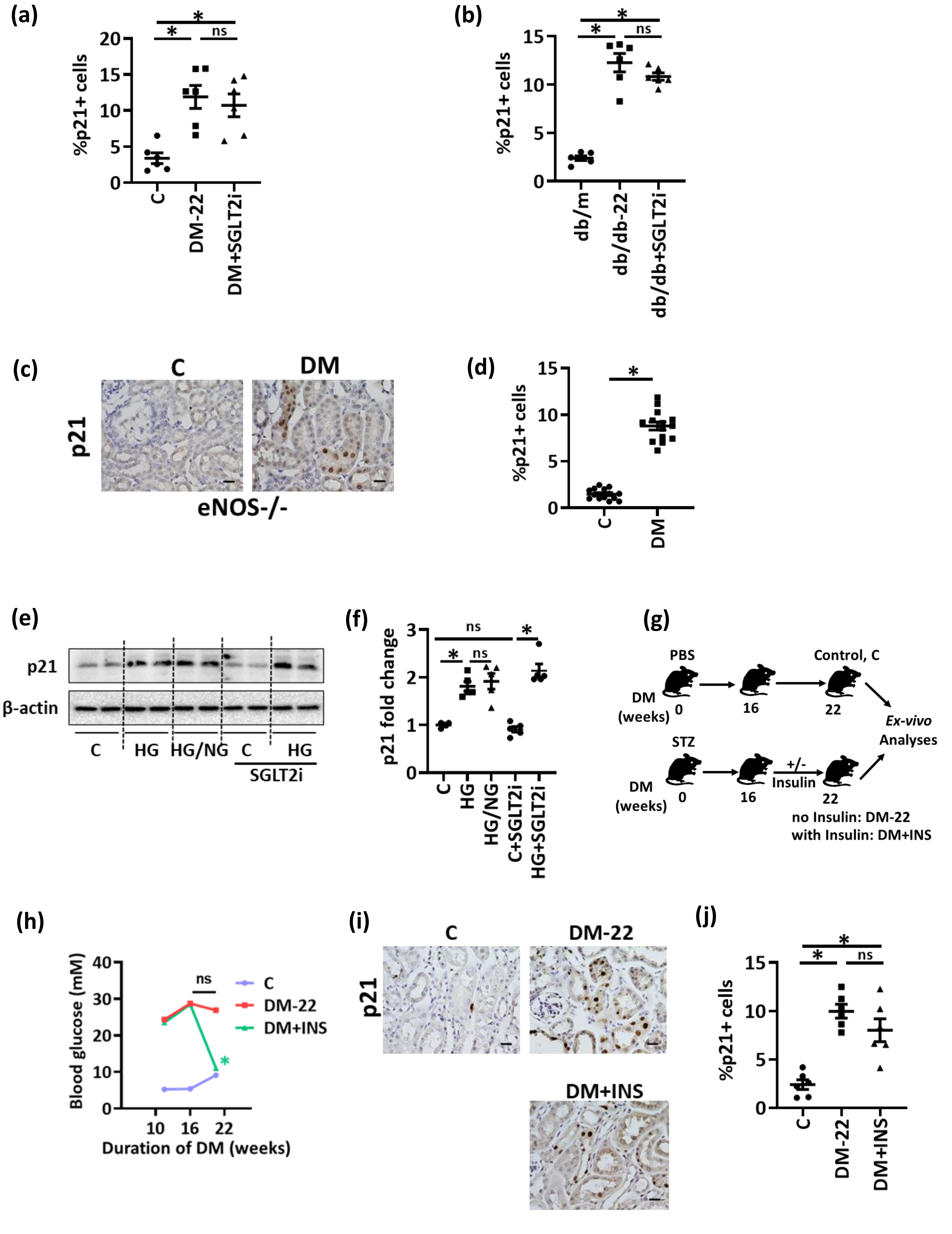

**Supp. Fig. 2: Sustained p21 expression despite reduced blood glucose levels in alternative experimental DKD in vivo and in vitro models**

**a,b)** Dot plots showing percentage of cells staining positive for p21 in vivo (T1DM, a; T2DM model, b) despite reducing glucose levels (DM+SGLT2i, a; db/db+SGLT2i, b) as compared to mice with persistently elevated glucose levels (DM-22, a; db/db, b) or normoglycemic controls (C, a; db/m, b). Dot plots reflecting mean±SEM of at least 6 mice per group; ANOVA, *P<0.05; ns: non-significant.

**c,d)** Exemplary histological images of p21 (c) in non-diabetic (C) or diabetic (T1DM) eNOS-/- mice (immunohistochemistry, p21 detected by HRP-DAB-reaction, brown; hematoxylin counterstain) and dot plot showing percentage of cells staining positive for p21 (d). Dot plots reflecting mean±SEM of at least 6 mice per group; ANOVA, *P<0.05.

**e,f)** Exemplary immunoblot of p21 (e; β-actin: loading control) in BUMPT cells and dot plot summarizing results (f). Experimental conditions: control with continuously normal glucose (C, 5 mM glucose), continuously high glucose (HG, 25 mM, 48 h), high glucose for 24 h followed by normal glucose (NG, 5 mM glucose) for 24 h, control with SGLT2i (Dapagliflozin®, 2 µM, C+SGLT2i), or continuously high glucose with SGLT2i (Dapagliflozin®, 2 µM, added in the second 24h, HG+SGLT2i). Dot plot (f) reflecting mean±SEM of at least 3 independent experiments; ANOVA, *P<0.05; ns: non-significant.

**g)** Experimental scheme showing non-diabetic control (C) or diabetic (T1DM) mice without (DM-22) or with intervention to reduce blood glucose levels by insulin (DM+INS). Mice were age matched and insulin treatment was started after 16 weeks of persistent hyperglycemia for further 6 weeks.

**h)** Average blood glucose levels in experimental groups (as described in g) after 10 or 16 weeks of hyperglycemia and at 22 weeks. Line graphs reflecting mean±SEM of at least 6 mice per group; ANOVA, *P<0.05.

**i,j)** Exemplary images of p21 (i, immunohistochemistry, p21 detected by HRP-DAB-reaction, brown; hematoxylin counterstain) and dot plot showing percentage of cells staining positive for p21 (j) in experimental groups (as described in g). Dot plot reflecting mean±SEM of at least 6 mice per group; ANOVA, *P<0.05; ns: non-significant.

#### Supp. Fig. 3 (related to Fig. 1)

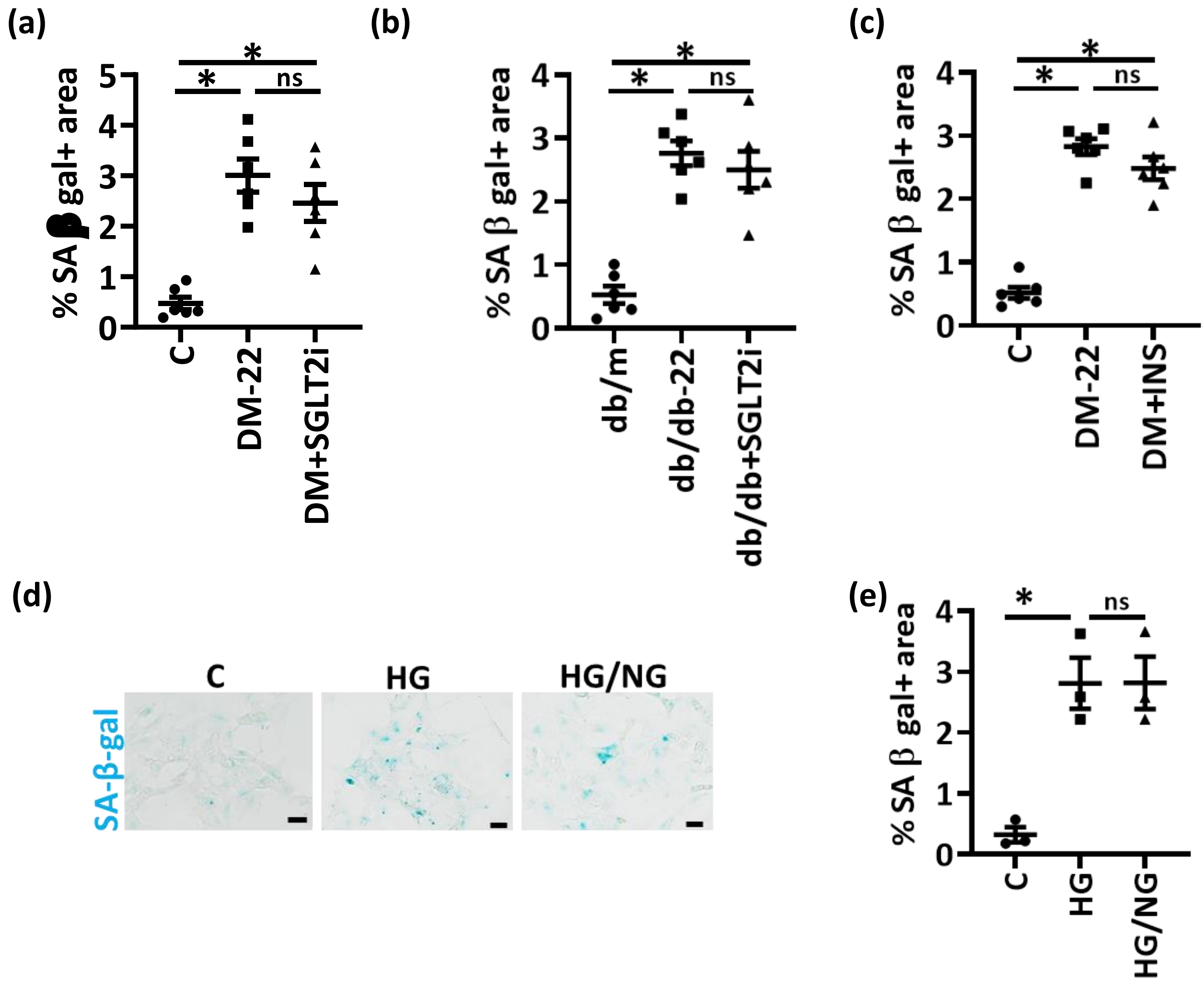


**Supp. fig. 3: Sustained renal tubular senescnece despite reduced blood glucose levels in experimental DKD and in renal cells *in vitro***

**a-c)** Dot plots showing percentage of SA-β-gal. positive area in vivo (T1DM model, a and c; T2DM model, b) despite reducing glucose levels (DM+SGLT2i, a; db/db+SGLT2i, b; or DM+INS, c) as compared to mice with persistently elevated glucose levels (DM-22, a and c; db/db, b) or normoglycemic controls (C, a and c; db/m, b). Dot plots reflecting mean±SEM of at least 6 mice per group; ANOVA, *P<0.05; ns: non-significant.

**d,e)** Exemplary images of SA-β-gal. stain (senescence associated β-galactosidase, blue, d) and dot plot showing percentage of SA-β-gal. positive area (e) in HEK-293 cells. Experimental conditions: control with continuously normal glucose (C, 5 mM glucose), continuously high glucose (HG, 25 mM, 48 h) or high glucose for 24 h followed by normal glucose (NG, 5 mM glucose) for 24 h. Dot plot reflecting mean±SEM of at least 3 independent experiments; ANOVA, *P<0.05; ns: non-significant.

#### Supp. Fig. 4 (related to Fig. 3)

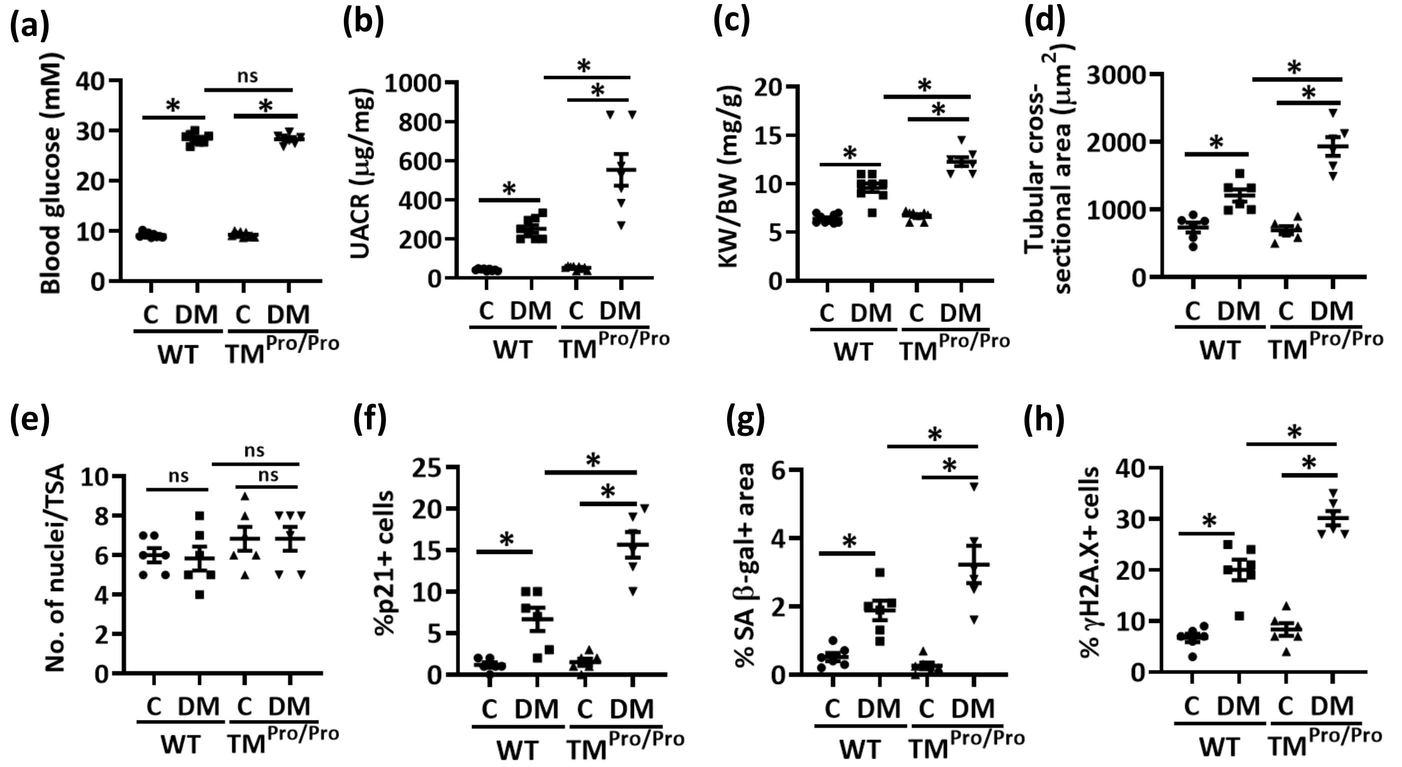


**Supp. Fig. 4: Loss of thrombomodulin-dependent protein C activation enhances tubular senescence and p21 expression**

**a-c)** Dot plots summarizing average blood glucose levels (a), albuminuria (urinary albumin-creatinine ratio, µg albumin/mg creatinine; UACR, b) and adjusted kidney weight (mg kidney weight /g body weight; KW/BW, c) in non-diabetic (C) or diabetic (DM) wild-type (Wt) and $TM^{Pro/Pro}$ mice. Dot plot reflecting mean ± SEM of at least 6 mice per group; ANOVA, *P<0.05; ns: non-significant.

**d-h)** Dot plots summarizing tubular hypertrophy (tubular cross-sectional area, TSA, d; and number of nuclei per TSA, e) and tubular cell senescence (p21 positive cells, f; SA-β-gal positive area, g; and γH2A.X positive cells, h) in experimental groups (as described in a). The number of nuclei per tubular cross-sectional area (TSA) is not different among experimental groups (e). Dot plots reflecting mean ± SEM of at least 6 mice per group. ANOVA; *P<0.05; ns: non-significant.

#### Supp. Fig. 5 (related to Fig. 4)

**(a)** **(b)**

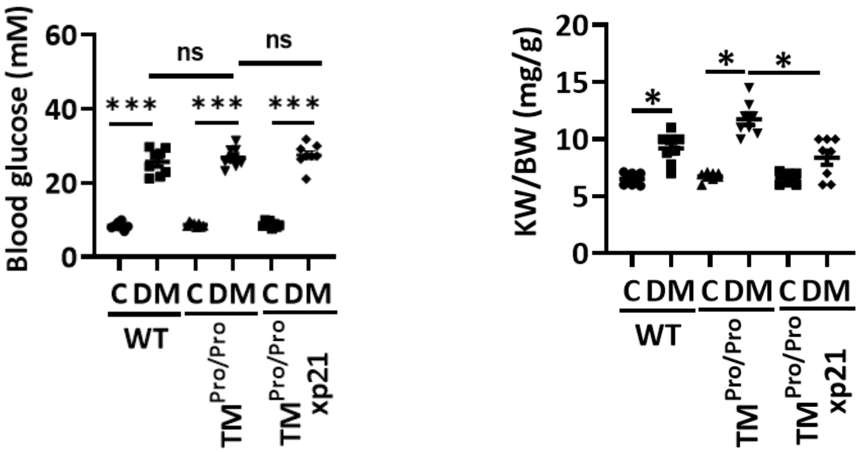


**Supp. Fig. 5: p21 promotes renal tubular senescence in aPC-deficient mice**

Dot plots summarizing average blood glucose levels (a) and adjusted kidney weight (mg kidney weight /g body weight; KW/BW, b) in non-diabetic (C) or diabetic (DM) wild-type (Wt), TM$^{Pro/Pro}$, or TM$^{Pro/Pro}$ x p21$^{-/-}$ mice. Dot plot reflecting mean±SEM of at least 6 mice per group; ANOVA, *P<0.05; ns: non-significant.

**Supp. Fig. 6 (related to Fig. 6)**

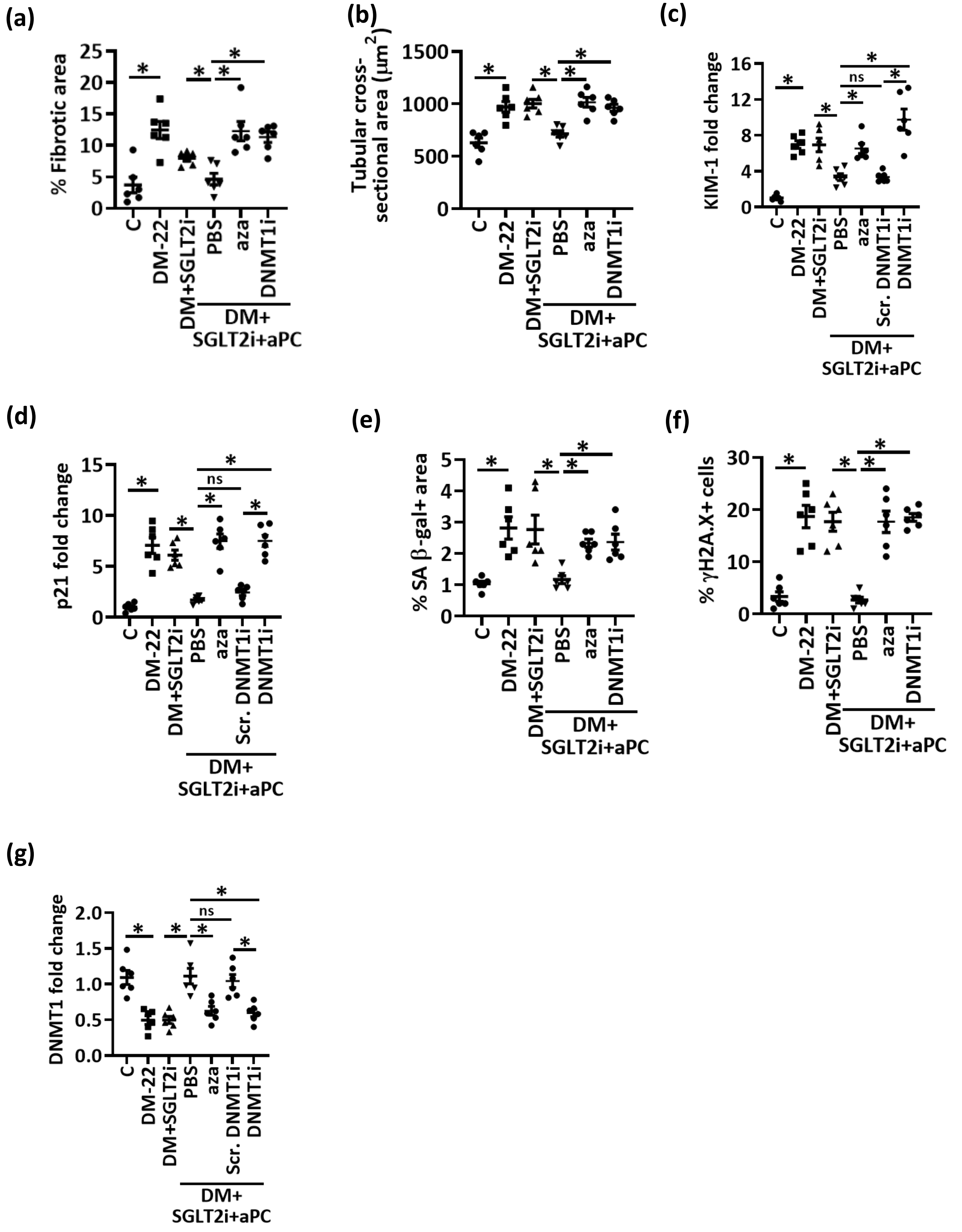

**Supp. Fig. 6: aPC reverses epigenetically sustained renal p21 expression and tubular senescence**

Dot plots summarizing tubulointerstitital damage (fibrotic area, a; tubular cross-sectional area, b), renal KIM-1 (c) and p21 (d) protein expression, tubular cell senescence (SA-β-gal positive area, e; γH2A.X positive cells, f) and renal DNMT1 protein expression (g) in non-diabetic (C) or diabetic (DM-22) mice. Subgroups of diabetic mice were treated with SGLT2i (DM+SGLT2i), SGLT2i with aPC and PBS (DM+SGLT2i+aPC+PBS), SGLT2i with aPC and 5-aza-2'-deoxycytidine (DM+SGLT2i+aPC+aza), SGLT2i with aPC and DNMT1 vivo morpholino (DM+SGLT2i+aPC+DMNTi), or SGLT2i with aPC and scrambled vivo morpholino (DM+ SGLT2i+aPC+Scr.DMNTi). Dot plots reflecting mean±SEM of at least 6 mice per group. ANOVA; **P*<0.05; ns: no significance.

### Supp. Fig. 7 (related Fig. 7)

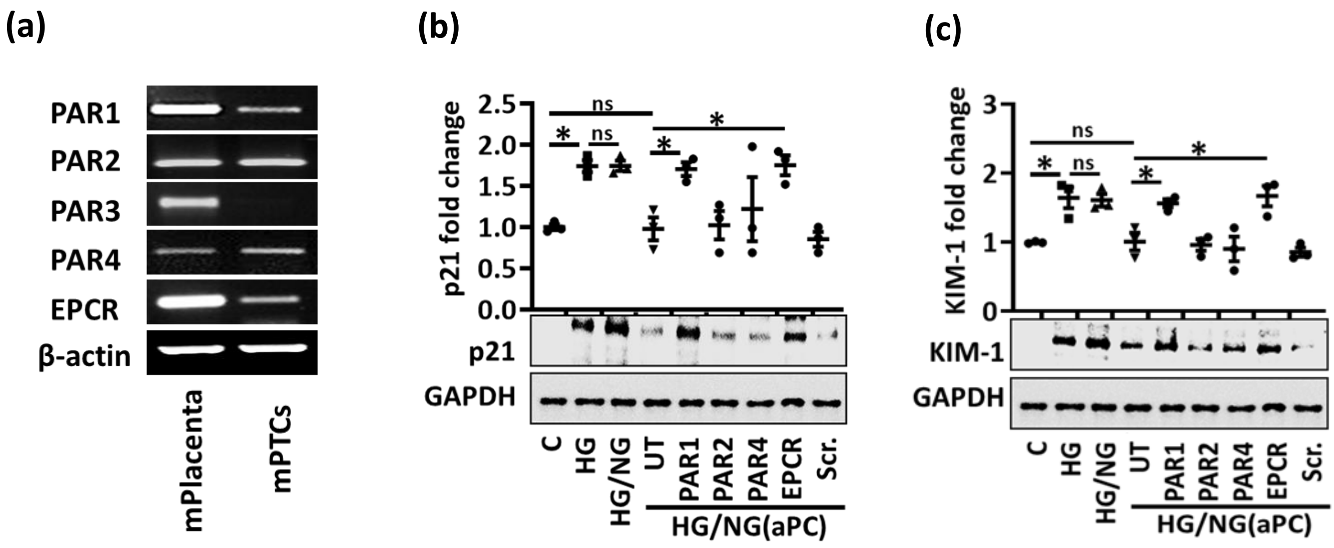


**Supp. Fig. 7: aPC suppresses glucose induced tubular cell p21 expression via PAR1 and EPCR**

**a)** Expression of PARs and EPCR in mouse proximal tubular cells (PTC; a; representative images of semiquantitative RT-PCR, mouse placenta as positive control).

**b,c)** aPC reverses sustained p21 (b) and KIM1 (c) expression in PTCs (HG/NG(aPC)). This effect is abolished upon PAR1 or EPCR knock down. Experimental conditions: control with continuously normal glucose (C, 5 mM glucose), continuously high glucose (HG, 25 mM, 48 h), or high glucose for 24 h followed by normal glucose (NG, 5 mM glucose) for 24 h without (HG/NG) or with aPC (20 nM) upon returning cells to normal glucose (HG/NG(aPC)). PAR1: PAR1 knock down; PAR2: PAR2 knock down; PAR4: PAR4 knock down; EPCR: EPCR knock down; Scr: non-specific, scrambled knock down construct. Exemplary immunoblots (bottom) and dot plots reflecting results (top). Dot plots reflecting mean±SEM of at least 3 independent experiments (b,c). ANOVA; **P*<0.05 comparing HG, HG/NG, and HG/NG(aPC) to C and knock down cells (PAR1, PAR2, PAR4, EPCR, Scrambled, Scr) to HG/NG(aPC); ns: non-significant.

#### Supp. Fig. 8 (related to Fig. 7)

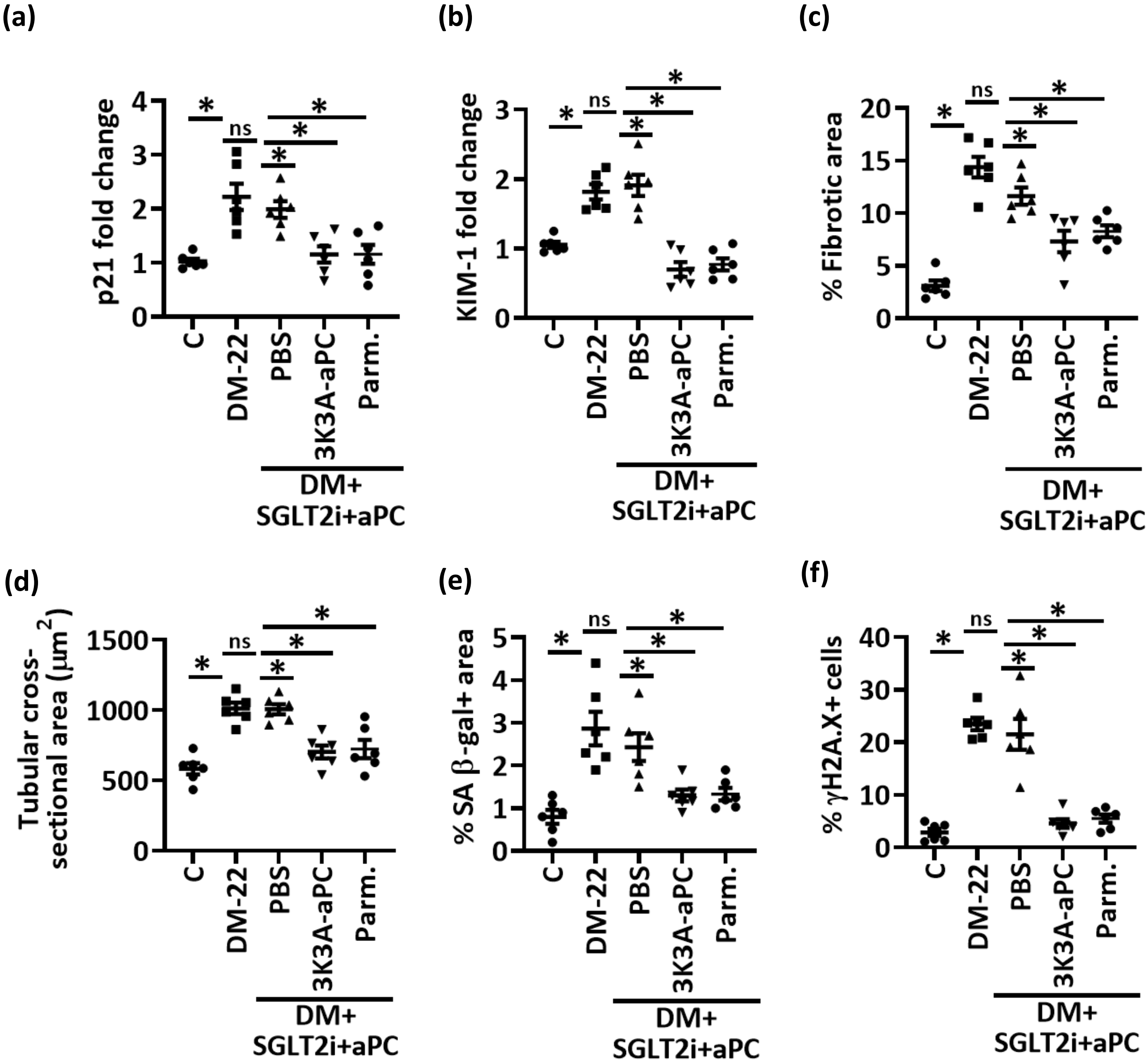


**Supp. Fig. 8: aPC reverses epigenetically sustained renal p21 expression and tubular senescence independent of its anticoagulant function**

**a,b)** Dot plots summarizing renal p21 (a) and KIM-1 (b) protein expression in non-diabetic (C) or diabetic (DM-22) mice. Subgroups of diabetic mice were treated with SGLT2i (DM+SGLT2i+PBS), SGLT2i with 3K3A-aPC (DM+SGLT2i+aPC+3K3A-aPC) or SGLT2i with parmodulin-2 (DM+SGLT2i+Parm.). Dot plots reflecting mean±SEM of at least 6 mice per group. ANOVA; *$P<0.05$; ns: non-significant.

**c-f)** Dot plots summarizing tubulointerstitital damage (fibrotic area, c; tubular cross-sectional area, d) and tubular cell senescence (SA-β-gal positive area, e; γH2A.X positive cells, f) in experimental groups (as described in a). Dot plots reflecting mean±SEM of at least 6 mice per group. ANOVA; *$P<0.05$; ns: non- significant.
